# Mechanism of the allosteric activation of the ClpP protease machinery by substrates and active-site inhibitors

**DOI:** 10.1101/578260

**Authors:** Jan Felix, Katharina Weinhäupl, Christophe Chipot, François Dehez, Audrey Hessel, Diego F. Gauto, Cecile Morlot, Olga Abian, Irina Gutsche, Adrian Velazquez-Campoy, Paul Schanda, Hugo Fraga

## Abstract

Coordinated conformational transitions in oligomeric enzymatic complexes modulate function in response to substrates and play a crucial role in enzyme inhibition and activation. ClpP protease is a tetradecameric complex that has emerged as a drug target against multiple pathogenic bacteria. During drug development efforts, the activation of different ClpPs by inhibitors was independently reported, but so far, no rationale for inhibitor-induced activation has been proposed. Using an integrated approach that included X-ray crystallography, solid-and solution-state NMR, MD simulations and ITC we show that the proteasome-inhibitor bortezomib binds to the ClpP active site serine mimicking a peptide substrate and induces the concerted allosteric activation of the complex. The bortezomib activated conformation also displays a higher affinity for its cognate unfoldase ClpX. We propose a universal allosteric mechanism where substrate binding to a single subunit locks ClpP into an active conformation optimized for chaperone association as well as protein processive degradation.

## Introduction

Among the most exciting antibacterial drug targets to emerge in the past decade is the caseinolytic protease (ClpP)(*1*). While mostly studied in *E. coli* (*Ec*ClpP), ClpP is present in a wide range of bacteria, as well as in mitochondria and chloroplasts. Functionally, ClpP can be defined as a cylindrical canonical serine protease composed of two stacked rings each formed by 7 subunits that encompass a large (300 kDa) proteolytic core (*2*). This architecture restricts access to the 14 proteolytic active sites located inside the barrel and is key for ClpP function. Although ClpP is able to rapidly hydrolyze peptides, the degradation of large proteins requires the presence of an AAA+ ATPase complex, such as ClpA or ClpX in *E. coli* or ClpC in other species. These unfoldases provide substrate specificity by recognizing, unfolding and translocating their globular protein substrates, but also by opening the apical gate region of ClpP, which is otherwise blocked by flexible N-terminal tails.

ClpP protease and its associated chaperones are essential for the survival or virulence of several bacteria including pathogens such as *Mycobacterium tuberculosis* (*Mtb*) (*3*), and *Staphylococcus aureus* (*Sa*)(*4*) and *Listeria monocytogenes* (*Lm*)(*5*). In addition, human mitochondrial ClpP was recently associated to human acute myeloid Leukemia and obesity (*6, 7*). Recently, ClpP from *Plasmodium falciparum* (*Pf*ClpP), the parasite responsible for malaria infection, has also been proposed as a promising drug target for the control of this disease (*8*).

The fact that ClpP is a suitable Achilles heel for these organisms is further reinforced by the recent discovery of natural antibiotics that target ClpP or its co-chaperones. Indeed, efficient bacterial killing could be accomplished either by opening the axial pore of ClpP and activating unregulated cell proteolysis(*9*), by blocking ClpP-ATPase interaction (*10*), or by specifically inhibiting its chaperones, ClpX (*4*) or ClpC1 (*11*).

Despite recent advances in the field, the study and development of drugs targeting ClpP has been complicated by the fact that ClpPs from different species show significant functional differences. For example, *Mtb* contains two clpP genes, clpP1 and clpP2, both of which are essential for viability and infectivity. Although both genes encode serine proteases that form heptameric rings, first attempts to express and characterize isolated *Mtb*ClpP1 and *Mtb*ClpP2 in *E. coli* yielded complexes that lacked proteolytic activity. The study of *Mtb*ClpP1P2 was not possible until the finding that a small activator, derived from classical serine protease inhibitor peptide aldehydes (*3*), was required to promote the formation of an active complex composed of one heptameric ring from ClpP1 and another from ClpP2 (*3*). Curiously, the small activators were found at the protein active site and it is not clear how their presence at the active site can promote activation rather than classic competitive inhibition (*12, 13*). Furthermore, several structures obtained by X-ray crystallography and solution NMR studies have suggested that ClpPs, while keeping the tetradecamer organization, can adopt several conformations. Currently, at least 3 conformations have been reported: an extended (catalytically active), a compressed (catalytically inactive), and a compact state (catalytically inactive) (*14*-*16*). In addition, while some ClpPs are inherently active, others are purified as non-active forms. Here we report our studies with ClpP from *Thermus thermophilus* (*Tt*ClpP) which is, as described for other ClpPs, purified in an inactive conformation, but can be *in vitro* activated by small molecules. Using an integrated approach including multiple biochemical assays, X-ray crystallography, magic-angle-spinning (MAS) NMR and solution-state NMR, isothermal titration calorimetry (ITC) and molecular-dynamics (MD) simulations, we show that bortezomib, a boronic acid previously identified as an inhibitor of *Mtb*ClpP1P2, induces a concerted conformational change in ClpP resulting in complex activation. Based on our data and previous results we propose a universal mechanism for the activation of ClpP.

## Results

### Bortezomib in low concentrations activates *Tt*ClpP peptidase activity

Although cellular protein quality control must be particularly challenging for organisms living at high temperatures, little has been described concerning ClpP from extremophiles. *Thermus thermophilus* (*Tt*) is a Gram negative eubacterium used in a wide range of biotechnological applications, and as a model organism for genetic manipulation, structural genomics, and systems biology. The bacterium has an optimal growth temperature of about 65 °C. We obtained *Thermus thermophilus* ClpP (*Tt*ClpP) by recombinant expression, and purified it as a canonical tetradecamer, as monitored by size exclusion chromatography. This tetradecamer could be reorganized into smaller species when ammonium sulfate (300 mM) was included in the elution buffer, likely as a result of the disruption of inter-ring salt bridge contacts (Fig. S1A). As anticipated, *Tt*ClpP displayed enhanced thermostability compared to previously studied ClpPs. Using the *Tt*ClpP monomer’s single tryptophan (14 per tetradecamer) as a probe, the T_*M*_ of ClpP was determined as 84 **°**C (Fig. S1B), 30 **°**C above the value reported for *S. aureus* ClpP (*Sa*ClpP)(*17*). The increased stability is also reflected by an elevated optimal catalytic temperature. When we measured the degradation of the green-fluorescent protein GFPsrrA by *Tt*ClpP in association with the cognate *Tt*ClpX ATPase, we found maximal rates at 60 **°**C (the highest temperature we could achieve with the fluorimeter equipment; Fig. S2A, S2B). However, although *Tt*ClpP is competent to degrade folded substrates in association with *Tt*ClpX, it is a rather inefficient peptidase even at 60°C compared to *Escherichia coli* ClpP (*Ec*ClpP) at 37**°**C it has a 200-fold lower specific activity in degradation experiments with the short tripeptide substrate PKMamc (Proline-Lysine-Methionine-7-amino-4-methylcoumarin, at a working concentration of 100 µM).

During attempts to identify inhibitors for *Tt*ClpP that could be used for further structural biology studies, we observed that bortezomib did not inhibit, but actually stimulated *Tt*ClpP peptidase activity at concentrations from 3 to 100 µM (Fig. 1A and Fig. S3A). Bortezomib is a N-protected dipeptide with a boronic acid instead of a carboxylic acid that forms adducts with activated threonines or serines (*18*). It was developed as a proteasome inhibitor targeting specifically the chymotrypsin-like sites of this complex with nanomolar affinity, whereas displaying limited reactivity to the other active sites of the complex. We selected this molecule because it was recently identified in an *in vivo* cell screen as an inhibitor of GFPssrA degradation by the *Mtb*ClpXP1P2 complex and was proposed as a potential anti-tuberculosis candidate (*19*). The bortezomib-dependence of the *Tt*ClpP peptidase activity displays a bell shape with maximum activation at 12.5 µM (about 10-20 fold activation), while higher drug concentrations reduced the enzymatic activity (Fig. 1A). A time lag phase for the activation of *Tt*ClpP was consistently observed and likely resulted from the necessary fluorimeter temperature equilibrium since the assays were executed at 45-60 **°**C. Bortezomib activation derives from intrinsic properties of *Tt*ClpP and when tested against *Ec*ClpP peptidase activity it behaved as a powerful inhibitor with an apparent IC50 of 1.6 µM (Fig. S3B).

**Figure 1.**
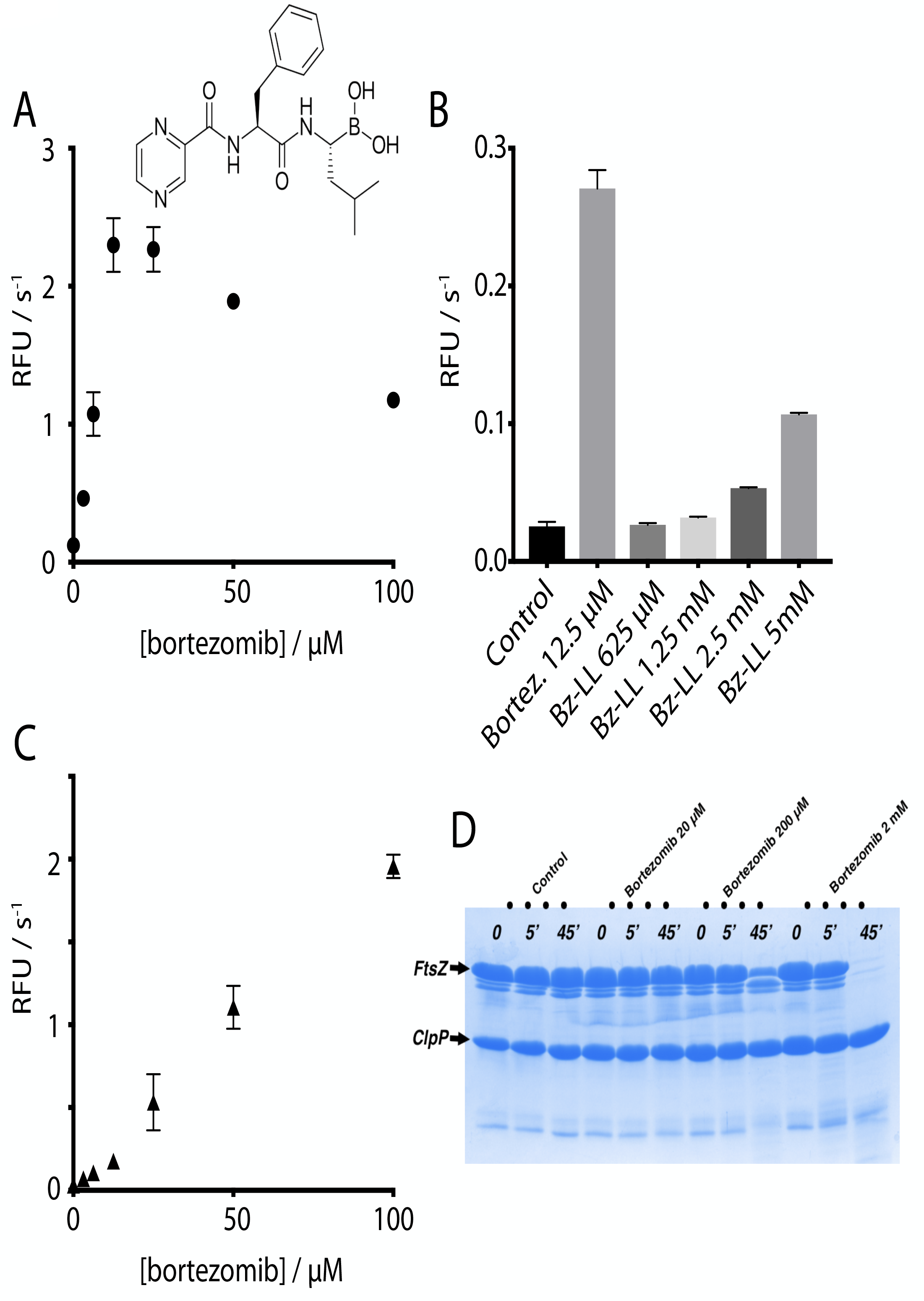
Bortezomib activates *Tt*ClpP for peptide and intrinsically-disordered protein degradation. A) *Tt*ClpP peptidase activity was measured with the substrate PKMamc (100 µM) in the presence of bortezomib at the indicated concentrations. As the peptide is cleaved, 7-Amino-4-methylcoumarin is released resulting in an increase in the measured fluorescence. Initial rates are plotted as a function of the bortezomib concentration. B) The activating effect of bortezomib is compared with the one observed with Bz-LL, a previously described activator of *Mtb*ClpP1P2. C) The degradation of the unfolded protein substrate FITC-casein by *Tt*ClpP was measured in the presence of the indicated bortezomib concentrations. Initial degradation rates following temperature equilibration were plotted as a function of the bortezomib concentration. D) *Tt*ClpP can degrade the intrinsically disordered *E.coli* FtsZ. The degradation of FtsZ by *Tt*ClpP was monitored by SDS-gel electrophoresis. While no degradation was observed with apo*Tt*ClpP, degradation of FtsZ was observed in the presence of 200 µM and 2 mM bortezomib.

Although paradoxical, the activation of some ClpP complexes by inhibitors has already been reported. Akopian and colleagues have shown that *Mtb*ClpP complex (*Mtb*ClpP1P2), otherwise inactive, is activated by small peptide aldehydes (*3*). In fact, the most potent of these activators in *Mtb*, a N-terminally blocked dileucine, (Bz-LL) was also capable, albeit with lower efficiency, to activate *Tt*ClpP (Fig. 1B). Vice versa, bortezomib could also activate the peptidase acti*vity* of the *Mtb*ClpP1P2 complex (Dr Tatos Akopian, Prof Alfred Goldberg, Harvard School of Public Health, personal communication, 2015). We applied a set of biochemical, biophysical and structural studies in order to understand the mechanistic and functional foundation of this inhibitor-based activation.

### Bortezomib activates *Tt*ClpP unfolded protein degradation

After establishing the bortezomib-induced activation of *Tt*ClpP for degradation of small peptides, we investigated its potential to stimulate protein degradation. Contrary to small peptides which can independently diffuse into the proteolytic chamber, the basal degradation of proteins by ClpP is low because the flexible N-terminal tails of ClpP act as a barrier to protein entry into the chamber. Our assay employed fluorescein isothiocyanate-labeled casein (FITC-casein), a proline-rich protein lacking stable secondary structures, as the proteolytic target. Protease-catalyzed hydrolysis of FITC-casein leads to highly fluorescent dye-labeled peptides. Similarly to the peptidase activity, addition of bortezomib did not result in inhibition of FITC-casein degradation, but instead a striking activation was observed (Fig.1C and Fig.S4A). Furthermore, contrary to what was observed with the peptidase activity, concentration dependence of this effect was linear and no reduction of the activation was observed up to 100 µM (Fig. 1C). In addition to casein, we tested if bortezomib could also promote the degradation of other substrates. The bacterial GTPase FtsZ forms a cytokinetic ring at midcell, recruits the division machinery and coordinates membrane and peptidoglycan cell wall invagination. A FtsZ monomer contains an intrinsically disordered C-terminal linker and the degradation of FtsZ by ClpP activated by the natural antibiotic acyldepsipeptides (ADEPs) has been proposed to provoke growth inhibition (*20*). We therefore tested the effect of bortezomib ClpP-catalyzed proteolysis on FtsZ and found that when FtsZ was incubated with ClpP in the presence of bortezomib, even at low temperature for *Tt*ClpP (37 **°**C), we observed its degradation (Fig.1D)(*20*). It is important to note the substantial differences in *Tt*ClpP concentration used for the different techniques, with 10-fold more protease being used for the FtsZ protein degradation study compared to FITC-casein degradation, as a consequence of the lower sensitivity of SDS-page staining versus fluorescein fluorescence methods. These experiments establish that bortezomib acts as a *Tt*ClpP activator for the degradation of different disordered protein substrates or small peptides.

### ClpX enhances peptidase activity of TtClpP

Since ClpP acts in association with specific chaperones *in vivo* (ClpX, ClpA or ClpC), we proceeded by assessing the effect of *Tt*ClpX association on *Tt*ClpP peptidase activity. When we measured the degradation of PKMamc by *Tt*ClpP in the presence of *Tt*ClpX, we observed a marked increase in peptidase activity that was dependent on *Tt*ClpX concentration (Fig. 2A, 2B). Similarly to *Tt*ClpP activation by bortezomib, a lag phase was observed that could result either from temperature equilibrium or from the oligomerization of *Tt*ClpX hexamers from monomers in the presence of ATP.

**Figure 2.**
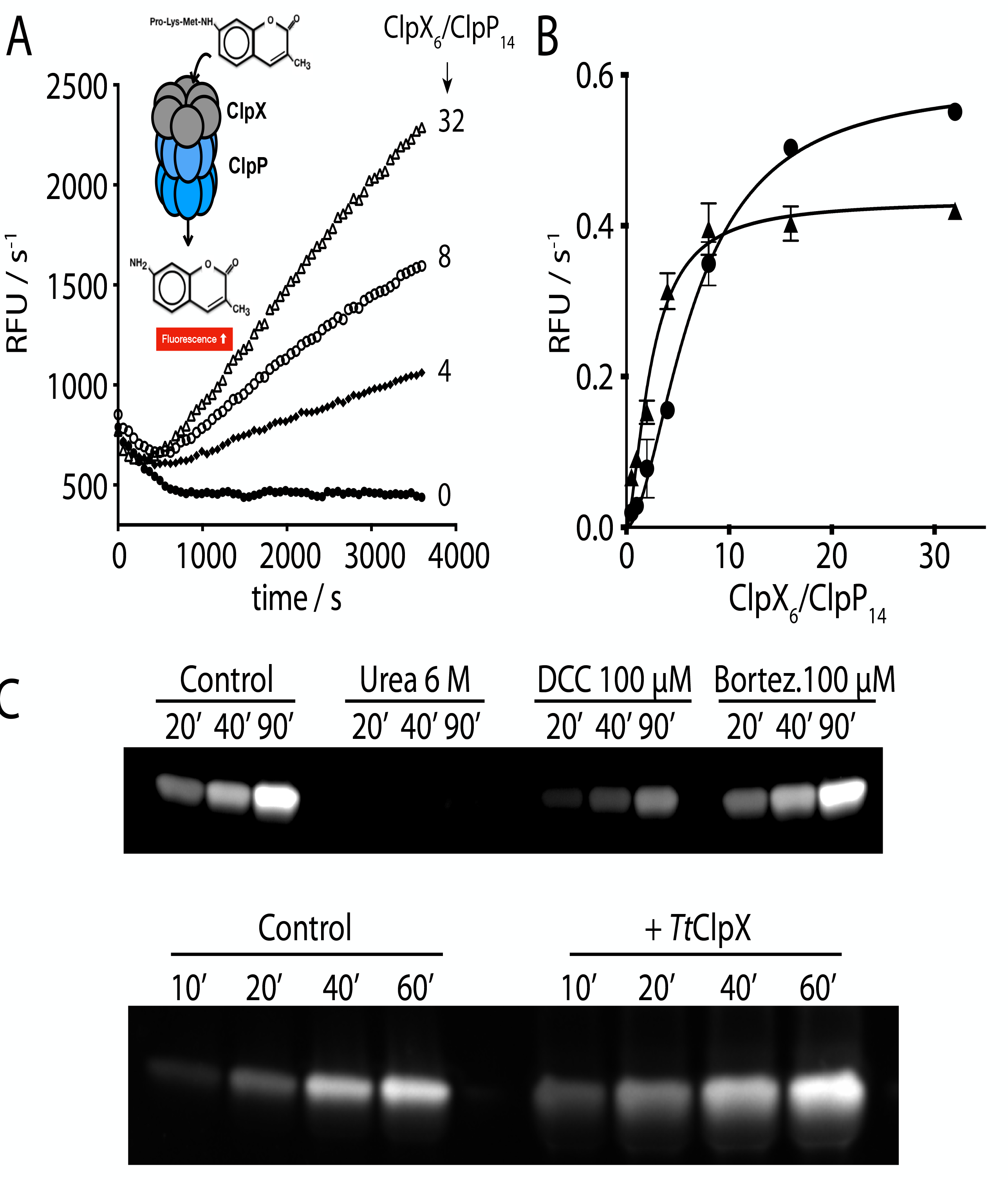
*Tt*ClpX activates *Tt*ClpP peptidase activity. A) *Tt*ClpP peptidase activity was measured in the presence of *Tt*ClpX at the indicated ClpX_6_/ClpP_14_ ratios. B) The activation of *Tt*ClpP peptidase activity in the absence or in the presence of bortezomib (10 µM) is plotted as a function of the molar ClpX_6_/ClpP_14_ ratio. Half maximal peptidase activity was obtained at 6.5 ± 0.5 and 2.4 ± 0.3 ClpX_6_/ClpP_14_ ratio for the apo and bortezomib loaded *Tt*ClpP respectively. C) Activation of the *Tt*ClpP active site by bortezomib and *Tt*ClpX, monitored by labeling with TAMRA-FP. TAMRA-FP was incubated with *Tt*ClpP and aliquots of the reaction mixture were removed at the indicated time points. Labelling of the active site serines is reflected by the increase in fluorescence observed.

Since bortezomib and ClpX are both able to activate *Tt*ClpP, we tested if the effects of bortezomib and ClpX were synergistic. Bortezomib (10 µM) did not induce a further activation, but acted instead as an inhibitor (Fig. S4B) in the presence of ClpX. However, a striking feature of the ClpX activation curve in the presence of bortezomib was a certain cooperative behavior, that is, less *Tt*ClpX concentration was required to achieve maximal activity, suggesting that bortezomib binding can modulate the affinity of ClpP for ClpX (Fig. 2B). Half maximal activation in the presence of bortezomib was achieved with a *Tt*ClpX_6_/ClpP_14_ ratio of 2.4, corresponding to 120 nM *Tt*ClpX_6_. This value for *Tt*ClpX-*Tt*ClpP association in the presence of bortezomib is in the same order of magnitude as the one reported for *E. coli* using an indirect ATPase assay(*21*).

The large activation of ClpP-catalyzed degradation of small peptides by ClpX was unexpected, since the auxiliary chaperones are required only for the unfolding and translocation of protein substrates into the ClpP proteolytic chamber, while small peptides can freely diffuse into the proteolytic chamber. Indeed, the hydrolysis rate of small peptides (didpetides) in *Ec*ClpP has been independently shown to be unaffected by ClpX association (*22, 23*). Activation by *Ec*ClpX is however essential for bigger peptides and proteins, and ClpX activation increases as a function of peptide molecular weight. ClpX binding has also been shown to stimulate active-site modification of ClpP by fluorophosphates, but only to a level expected from faster diffusion of the inhibitor into the ClpP chamber (*22*). Nonetheless, it was recently proposed that, in addition to the opening of the ClpP gate, chaperones can also allosterically activate the ClpP catalytic sites, by inducing a shift of ClpP from the inactive compressed state, in which the catalytic triad is distorted, to the extended state in which the catalytic residues are positioned for enzymatic action (*17*).

Since the small molecular weight of the substrate used in our assays is not an obstacle to its free diffusion into the proteolytic chamber (MW: 574 Da)(*22*), we tested if *Tt*ClpX binding to *Tt*ClpP could affect the protease active sites. In order to confirm the effect of co-chaperone or bortezomib on the active site, we used an assay based on TAMRA-FP (MW: 680 Da), a fluorophore-conjugated fluorophosphonate compound which specifically and covalently labels activated Ser-active sites with a fluorescent dye. TAMRA-FP can be used to quantify the reactivity of a given protease active site, for example if the catalytic triad is in the correct geometry for proper serine activation. The presence of an activated Ser can thus be detected using fluorescent gel imaging. We incubated *Tt*ClpP and TAMRA-FP with either urea, bortezomib, dichloroisocoumarin (DCC, an inhibitor of serine proteases), or *Tt*ClpX. While urea, by unfolding the complex, and DCC by irreversibly blocking the active site(*24*), reduced the amount of TAMRA bound to ClpP, ClpX association resulted in a significant increase in active site acetylation – a clear indication that ClpX association activates ClpP active site. Though bortezomib also leads to an activation of ClpP, we failed to observe any increase in TAMRA-FP labeling (Fig 2C). This could result from competition between the two inhibitors (bortezomib and TAMRA-FP), that is, we cannot exclude that TAMRA-FP binding also results in *Tt*ClpP activation. However, contrary to bortezomib, TAMRA-FP forms irreversible adducts with *Tt*ClpP, thereby preventing activity measurements.

### Bortezomib binds to the serine of ClpP active site

To understand bortezomib’s mechanism of action, we probed the bortezomib binding site using NMR spectroscopy. With a molecular weight of 300 kDa, NMR studies of *Tt*ClpP are challenging. In solution, the slow tumbling leads to severe line broadening, such that standard solution-NMR studies are not feasible. Essentially the only viable route for observing solution-NMR signals of proteins of the size of ClpP is the specific labeling of methyl groups in an otherwise deuterated background (*25*). Although being a very powerful approach for probing interactions and dynamics, methyl-directed NMR is able to observe only a subset of amino-acid types, and the sequence-specific assignment of NMR resonances is challenging, generally requiring multiple mutants.

MAS solid-state NMR does not suffer from this inherent size limitation, and is able to detect all amino acid types. We have used samples of *Tt*ClpP that were either obtained from sedimenting ClpP from solution into MAS NMR sample tubes (1.3 mm rotors for MAS) or obtained by addition of methyl-pentane-diol, an oft-used precipitant in crystallography. Spectra from these two preparations are very similar. We have used proton-detected three-and four-dimensional MAS NMR approaches (*26*) to obtain the sequence-specific resonance assignment for (^1^H^N^, ^15^N, ^13^Cα, ^13^CO) nuclei of 97 residues spread throughout the molecule, including the active site and the central helix αE which undergoes most changes in the compressed-to-extended transition. The N-terminal gate region was, however, not assigned, presumably due to its structural heterogeneity and dynamics which lead to inefficient NMR coherence transfers and line broadening. Fig. 3A shows the residue-wise secondary structure, obtained from these MAS NMR resonance assignments. The location of secondary structures matches well the information obtained from crystallography (see below), as expected.

**Figure 3.**
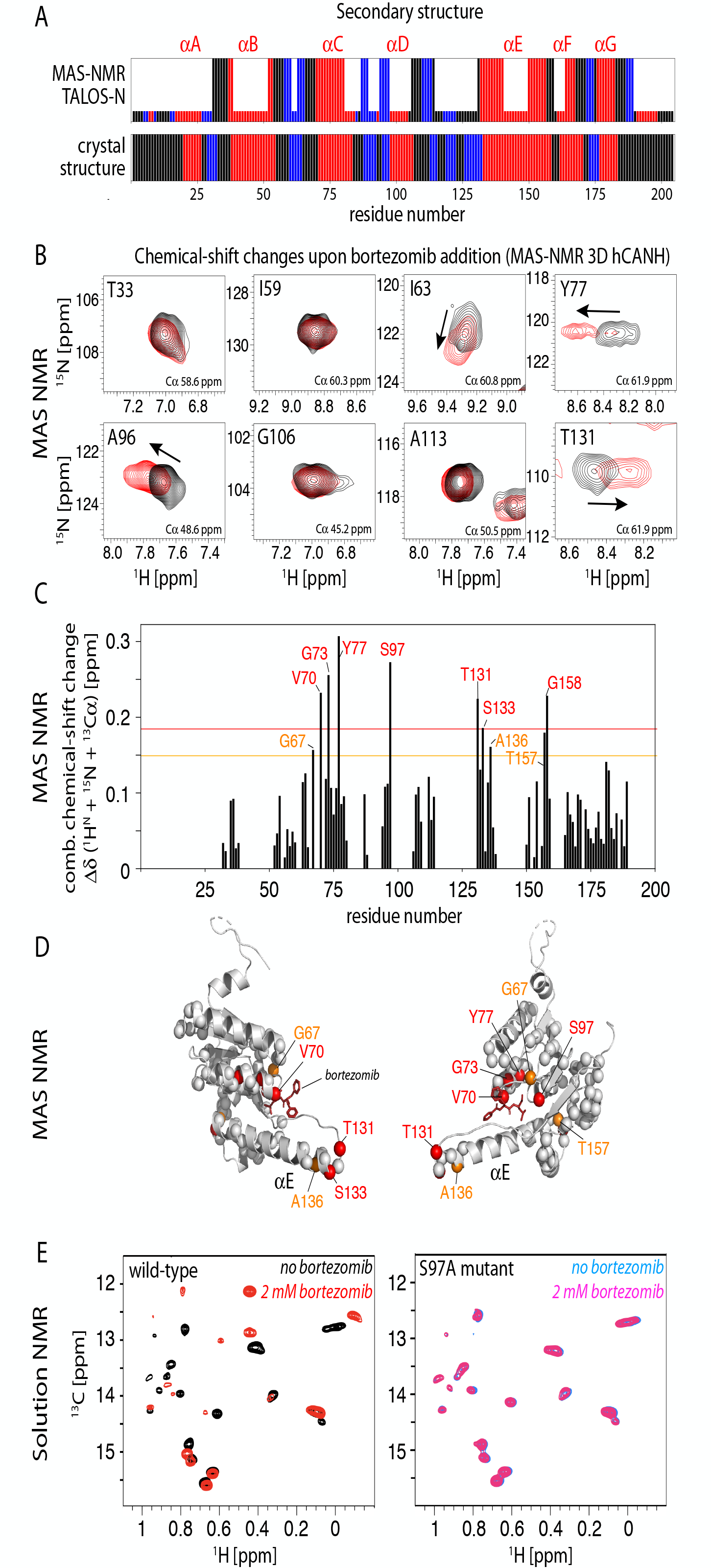
Bortezomib binding to TtClpP investigated by MAS NMR and solution-NMR. A) Secondary structures (loop: black, α-helix: red, β-strand: blue) as a function of sequence number obtained from the resonance assignments obtained by MAS NMR and analysis with TALOS-N(*45*) (tall bars). For residues for which no assignment has been obtained, the small bars are obtained by TALOS-N from a database approach. The lower panel shows the comparison with the secondary structure obtained from the crystal structure (see Figure 5). B) Zooms onto two-dimensional ^1^H-^15^N excerpts from 3D hCANH MAS NMR correlation spectra of *Tt*ClpP sedimented in the presence of 10 mM bortezomib (red) or without bortezomib (black). Chemical-shift changes are indicated with arrows. Note that the CSPs are relatively small, presumably due to the difficulty of saturating the binding site in a sedimented sample, given the high K_D_ value, resulting in incompletely occupied binding sites. C) Combined chemical-shift perturbations upon addition of bortezomib from the added peak shifts in ^1^H, ^13^Cα and ^15^N dimensions, obtained from the spectra shown in (B). D) Location of residues with significant chemical-shift perturbations mapped onto the X-ray structure of the *Tt*ClpP:bortezomib complex, obtained from the data shown in (C). E) Solution-state NMR spectra of wild-type (left) and S97A (right) *Tt*ClpP in the absence and presence of bortezomib, showing large spectra changes for the former due to binding, but no effect on the mutant.

Three-dimensional hCANH MAS-NMR spectra of sedimented *Tt*ClpP with and without bortezomib are shown in Fig. 3B and C. The observed changes are modest, which is likely linked to the relatively low affinity of bortezomib which makes it difficult to saturate the protein in the highly concentrated sedimented sample. Nonetheless, chemical-shift perturbations (CSP) are observed for a number of residues in the vicinity of the active site, including Val70, Gly73, Tyr77 and the catalytic Ser97, as well as residues in the neighboring helix αE (Fig. 3D). These data show that bortezomib interacts with the active site, and suggest that the interaction induces also small changes of structure and dynamics in the helix αE, which is involved in mediating the compressed-to-extended transition.

To independently confirm the direct involvement of the active site in bortezomib-binding in solution, we have performed additional NMR experiments, making use of perdeuterated, Ile-δ1-CH3 labelled *Tt*ClpP. Spectra of apo *Tt*ClpP, as well as in the presence of 2 mM bortezomib are shown in Fig. 3E (left panel). Although we did not attempt to assign the resonances of the 18 isoleucines per ClpP subunit, the clear differences in the spectra without/with bortezomib unambiguously shows the binding. In a control experiment performed with the catalytic S97A mutant, no spectral changes were observed, confirming that the bortezomib interaction requires the presence of the catalytic serine (Fig. 3E, right panel).

Taken together, MAS NMR and solution NMR provide direct evidence that bortezomib binds to the catalytic site. Chemical-shift perturbations outside the active site (in particular in helix αE) suggest that binding may lead to further structural changes, possibly linked to the observed *Tt*ClpP activation.

### Structural basis for bortezomib activation

In order to understand the structural details of the *Tt*ClpP-bortezomib interaction initially characterized by NMR, we used X-ray crystallography and determined the crystal structures of *Tt*ClpP in the absence and presence of bortezomib at a resolution of 1.95 Å and 2.7 Å, respectively (Fig. 4).

**Figure 4.**
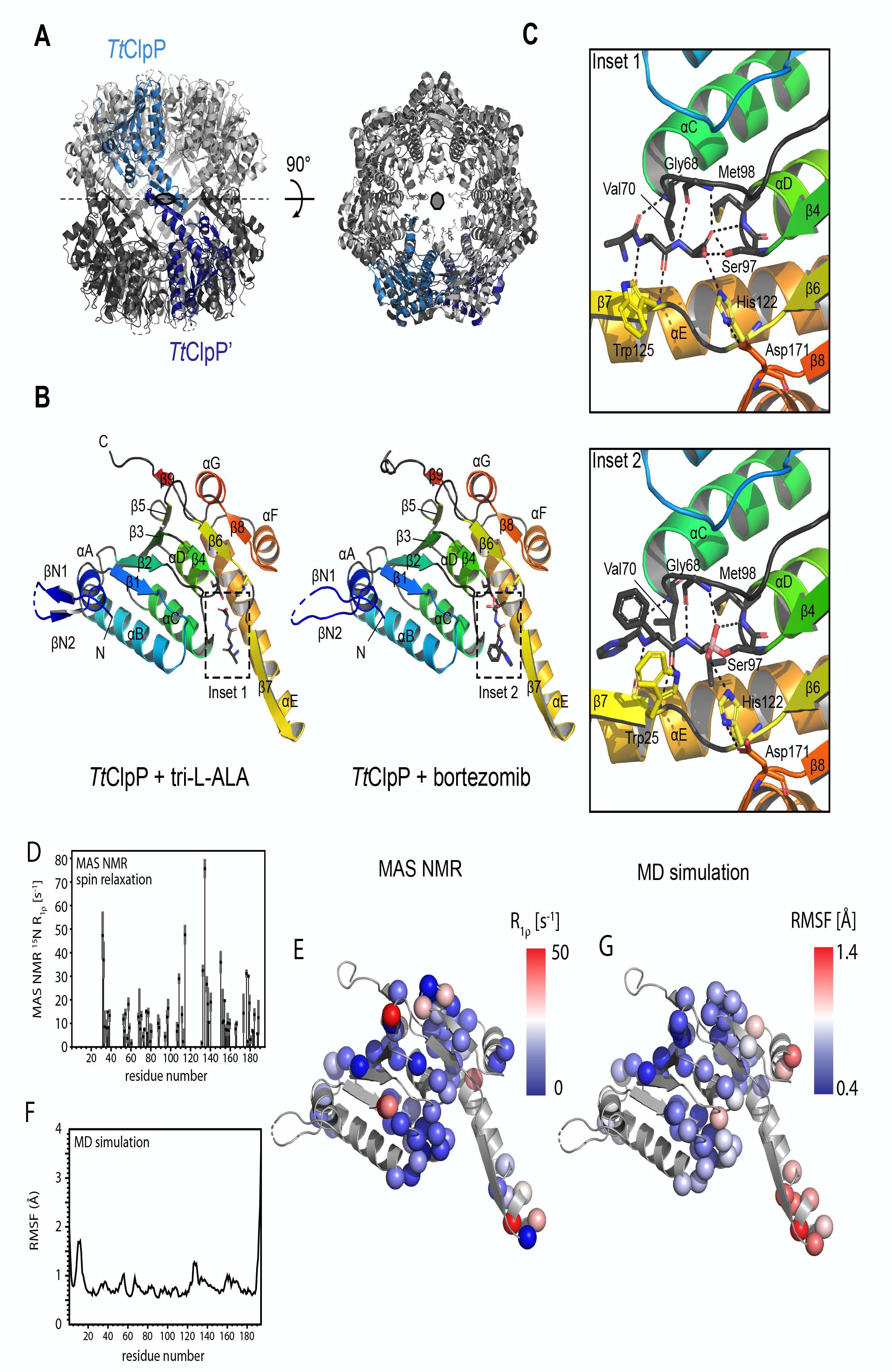
Structure and dynamics of *Tt*ClpP from X-ray crystallography, MAS NMR and MD simulations. A) Side and top views of the *Tt*ClpP 14-mer. One TtClpP monomer per heptameric ring (light and dark grey) is highlighted in light and dark blue respectively. B) Cartoon representation of the *Tt*ClpP monomer in peptide (left) and bortezomib-bound (right) forms. Helices are named by letters and strands are indicated by numbers. A zoom of the ligands present in the active sites (dashed boxes) are shown in C. C) Substrate binding pocket of *Tt*ClpP. The residues involved in the binding to the model peptide (inset 1) and bortezomib (inset 2) are shown as sticks. D,E) Residue-wise MAS NMR amide-^15^N R_1⍴_ relaxation rate constants. High R_1⍴_ rate constants point to enhanced ns-ms motions, and are found primarily in loop regions and in helix αE, as shown in panel C). F,G) MD-derived root-mean-square-fluctuations (RMSF) over the 1 µs-long MD trajectory of the assembled 14-mer ClpP.

The overall structure of *Tt*ClpP is essentially identical to the ‘extended’ state (*27*-*29*) of ClpP described for other species (r.m.s.d. to *Ec*ClpP tetradecamer [PDB id 6MT6] = 2.115 Å for 1451 aligned Cα atoms, r.m.s.d. to *Ec*ClpP monomer = 0.73 Å for 176 aligned Cα atoms), which is in in agreement with its high sequence similarity to ClpP orthologs (% identity to *Ec*ClpP = 58.25). The structure of one *Tt*ClpP monomer is presented in Fig. 4B, and displays an α/β fold composed of two domains: a head domain comprising six repeats of α/β units (helices A-D, F, G and strands 1-5, 7 and 8) and a handle domain (helix E and strand 6). The N-terminus of *Tt*ClpP forms a short anti-parallel β-hairpin that protrudes from the apical surface of the barrel. This N-terminal β-hairpin is constituted by residues 5-8 and 15-18 of *Tt*ClpP, with missing electron density for residues 9-14. While the overall monomer structure of ClpP is highly conserved among the structures solved to date, a high degree of variability exists in the conformation of the N-terminal regions. Several ClpP crystal structures display N-terminal β-hairpin structures of varying lengths and degrees of flexibility (*30*), of which some seem to be stabilized by crystal packing contacts (*16*, *30*, *31*). Solution NMR studies have demonstrated that the N-termini of ClpP are structurally heterogeneous and adopt multiple conformations (*32, 33*), which might explain the missing density near the end of the N-terminal β-hairpins in the *Tt*ClpP structure. Surprisingly, in the apo-*Tt*ClpP crystals, we consistently observe a continuous stretch of electron density near the active site serine (Ser97) of each *Tt*ClpP monomer, which based on its shape, suggests the presence of a tripeptide (Fig. 4C, Fig. S5). As no peptides have been added, the nature of this molecule was unknown, and we have built it in the electron density as a tri-L-alanine (see also omit and polder maps presented in Fig. S5). The presence of short peptides in *Helicobacter pylor*i ClpP structures resulting from proteolysis during crystallization has been previously described (*34*). While in that case the observed tri-and tetrapeptides are caused by proteolysis of added heptapeptides prior to crystallization, the origin of the tripeptides observed in our *Tt*ClpP crystals is unclear. Maldi-TOF measurements exclude the presence of these peptides in fresh purified TtClpP and we speculate that they are the result of auto-proteolysis of unfolded or partially unfolded *Tt*ClpP in the crystallization drops followed by the formation of *Tt*ClpP-peptide crystals. The serendipitous presence of peptides near the active site Ser97 provides some valuable insights on how *TtClpP* binds its substrates (Fig. 4C, insert 1). Apart from the active site Ser97, the terminal carboxy group of the peptide (residue P3) is further stabilized by hydrogen bonds with the Nε2 atom of His122 (part of the ClpP catalytic triad) and the backbone amide groups of Met98 and Gly68. The Nδ1 atom of His122 on its turn makes an additional hydrogen bond with Asp171, the third residue of the ClpP catalytic triad. The amino groups of peptide residues P2 and P3 are stabilized by backbone interactions with Gly68 and Trp125 respectively, while the carbonyl groups are stabilized by interactions with amino groups of Trp125 and Val70 (Fig. 4C, insert 1). The geometry and distances between the catalytic triad residues (Ser97, His122 and Asp171) are consistent with a functional *Tt*ClpP catalytic triad since their distance are in the range of the ones observed in classical serine proteases and active ClpP’s (*34*). Fig. 4B displays the 2.7 Å resolution structure of the bortezomib-bound state (see also Fig. S5). Similarly to the peptides present in the apo-*Tt*ClpP structure, bortezomib tightly fits between β-strand 6 and a small β*-*turn formed by Gly68 and Val70 thereby forming an antiparallel β*-*sheet. This conformation is characterized by a hydrogen network almost identical to the one observed in the *Tt*ClpP:peptide interaction (Fig. 4C, insert 2). Unlike the *Tt*ClpP:peptide complex, where Ser97 is forming hydrogen bonds with the terminal carboxy group of the peptide, bortezomib seems to be covalently linked to the active site Ser97 via its boron atom. Other significant differences are observed in the conformation of Trp125, which is displaced towards bortezomib and forms an aromatic interaction with the bortezomib phenylalanyl group. In addition, the nearby Thr131 residue, which in peptide bound *Tt*ClpP interacts with Arg170 of a neighboring ClpP monomer, now makes a hydrogen bond with Gln123 of an opposing ClpP monomer in the other heptameric ring. Despite the rearrangement of the Trp125 side chain, the catalytic triad geometry is again consistent with a fully active protease.

To probe the flexibility of *Tt*ClpP, we used MAS NMR spin-relaxation experiments and molecular dynamics simulations. Site-specific amide-^15^N R_1⍴_ experiments probe the local amplitudes and time scales of backbone motion, and are particularly sensitive to nanosecond-to-millisecond motions (*35*). Interestingly, the sites with highest ^15^N R_1⍴_ rate constants are found not only in loop regions (where flexibility is expected), but also at the tip of helix αE (Fig. 4D,E). One-µs-long MD simulations of the *Tt*ClpP tetradecamer in apo and peptide-bound states corroborate this observation, with a greater flexibility at the tip of the helix (Fig. 4F,G). Enhanced flexibility in this helix has been reported also in *Ec*ClpP by methyl-directed solution-state NMR (*15*). Due to the lack of reliable parametrization of the boronic acid group of bortezomib, we focused on simulating tripeptides bound to the active site. It is noteworthy that over the length of three independent simulations of ClpP that was initially loaded with tri-alanine in all active sites, only a fourth of the peptides remain associated to their designated active sites (Fig. S6), consistent with the idea that the substrates of *Tt*ClpP are necessarily weak binders, ought to be released rapidly to leave the catalytic sites ready for the next degradation reaction.

### Bortezomib induces the transition from a tense state to a relaxed state with higher affinity for the substrate

Having established the bortezomib-binding site on *Tt*ClpP by NMR and X-ray crystallography, we used ITC to thermodynamically characterize this interaction at different temperatures, from 25°C to 45°C. At 25 **°**C, far from the ideal catalytic temperature of *Tt*ClpP, only a moderate affinity and symmetric binding isotherm was observed. The binding enthalpy was negative and rather small (see Table I) indicating that binding was mainly entropy-driven. However, when the temperature was increased to 35 and 45 **°**C (values closer to the optimal catalytic temperature of this protein), significant changes were observed in the binding isotherm. The asymmetry of the curve and the appearance of two different phases in the binding isotherm along the titration, within the low ligand saturation region, were clear indicators for positive cooperativity (Fig. 5A). Because of the architecture of *Tt*ClpP, comprising two heptameric rings forming a tetradecameric assembly, ITC data was fitted to the Monod, Wyman and Changeux (MWC) model (Fig. 5B), which considers two intrinsically non-cooperative conformations, with low and high binding affinity, denoted as tense (T) and relaxed (R) state, respectively. In the MWC model, ligand binding shifts the equilibrium towards the high affinity conformation (R) in a concerted cooperative way (all subunits change their conformation simultaneously; Table I). This model has been previously implemented to other enzymatic systems, where activation at low ligand concentration and inactivation at high ligand concentration was also observed (*36*). Fitting of the ITC data revealed that in the absence of ligand 76% of *Tt*ClpP exists in the T state, which has lower affinity for bortezomib (K_d,T_ = K_T_^-1^ = 40 µM). Bortezomib binding induces a coordinated transition from the T to the R state with higher affinity for substrate (K_d,R_ = K_R_^-1^ = 15 µM). A model in which we assumed that there is no ligand binding to the subunits in T conformation (K_T_ =0, ΔH_T_ = 0) can be ruled out, based on significantly worse fitting (data not shown). The higher affinity of the ligand for the R conformation drives the conversion of T subunits into R subunits. In addition, the compulsory concerted conformational change within a given protein oligomer (all subunits in a given protein oligomer must undergo the conformational change at once, even if not all of them are occupied by ligand) will further promote the conversion of T subunits into R subunits, increasing the total fraction of R subunits. The Hill coefficient (maximal slope in the log(n_LB_/14-n_LB_) vs. log[ligand] plot) of *Tt*ClpP is 1.3, which is rather small for a tetradecameric protein, but was expected considering the small value for the conformational equilibrium constant γ.

**Table 1.**
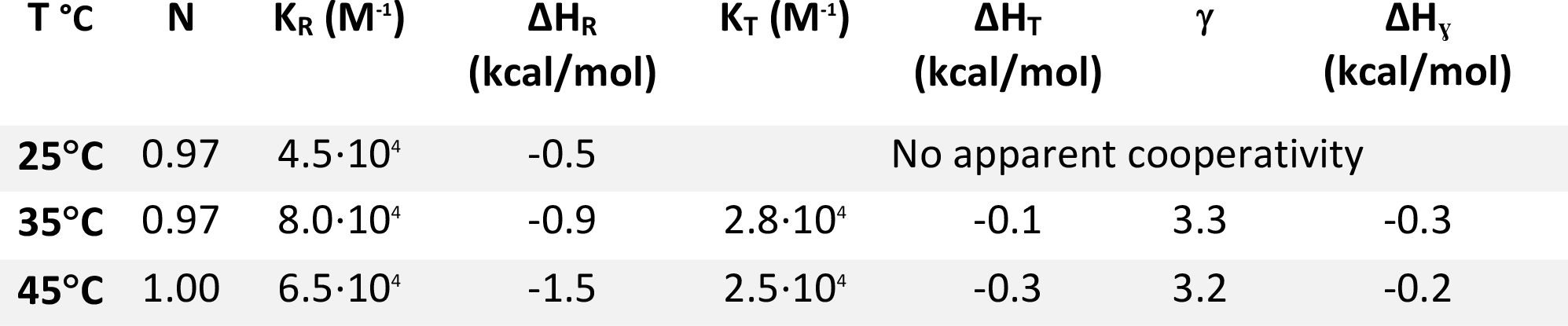
Thermodynamic parameters of bortezomib binding from ITC and fitting to the MWC model. K_R_ and K_T_ refer to the bortezomib associations constants for the relaxed and tense state. ΔH_R_, ΔH_T_ and ΔHγ refer to the binding enthalpy of the R state, T state and conformational enthalpy between R and T conformations. γ is the equilibrium constant between R and T conformations in the absence of ligand. N is the fraction of active or binding-competent protein. Relative errors in equilibrium constants are 30%/ Absolute error in enthalpies is 0.3 kcal/mol.

**Figure 5.**
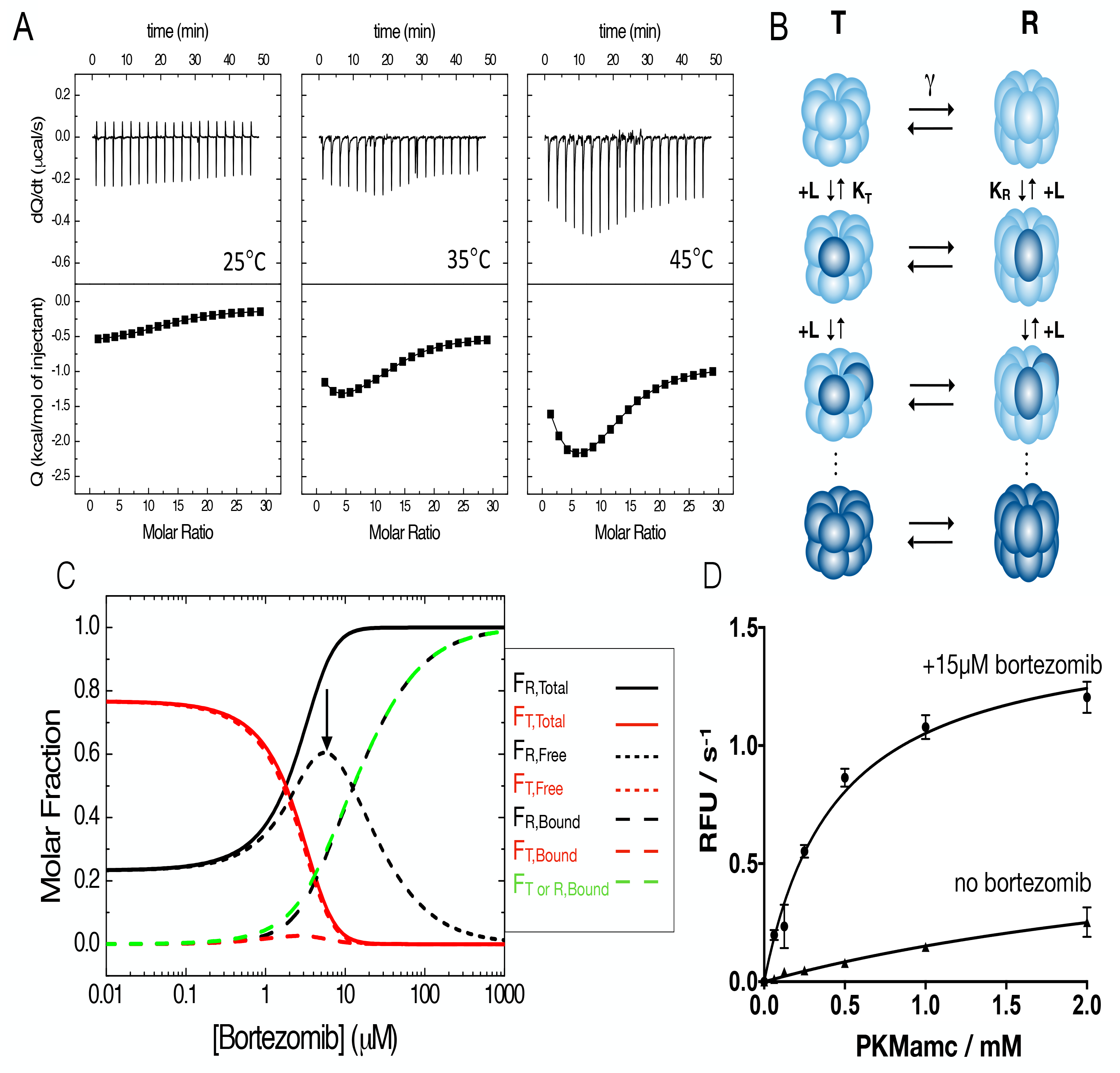
Cooperative bortezomib binding detected by ITC experiments. A) Calorimetric titrations for the interaction of ClpP (10 µM in the calorimetric cell) with bortezomib (1.4 mM in the injecting syringe) in Hepes pH 7.6 50 mM, NaCl 50 mM. Experiments were performed at three different temperatures: 25, 35 and 45 °C. Thermograms (thermal power as a function of time) are displayed in the upper plots, and binding isotherms (ligand-normalized heat effects per injection as a function of the molar ratio, [L]_T_/[P]_T_) are displayed in the lower plots. The binding isotherms were analyzed with the MWC model for ClpP, an oligomeric macromolecule consisting of 14 identical subunits, each one containing a single ligand binding site. Non-linear least squares regression analysis allows determining the binding parameters (see Table I): association constants for the R and T states (K_R_, K_T_), binding enthalpies to the R and T states (ΔH_R_, ΔH_T_), conformational equilibrium constant and conformational enthalpy change between states R and T (γ, ΔHγ), and fraction of active protein (N). B) MWC model for a 14-mer oligomeric protein. The protein can populate only two conformational states in equilibrium (with equilibrium constant γ): all subunits in R (relaxed, ellipsoidal shape) conformation or all subunits in T (tense, spherical shape) conformation. Subunits have a single ligand binding site, exhibiting the R conformation a higher binding affinity (K_R_ > K_T_). Ligand binding occurs through an independent (non-cooperative) fashion within an oligomer (ligand-free subunits in light blue, ligand-bound subunits in dark blue). T conformation is favored at low ligand concentration; the higher ligand binding affinity for R conformation promote a highly cooperative compulsory concerted conformational change driven by ligand binding and involving all subunits within an oligomer, displacing the equilibrium towards the R conformation. C) Molar fraction of the different protein species (total, ligand-free and ligand-bound R and T conformations) as a function of ligand concentration: total fraction of subunits in R conformation (continuous black line), total fraction of subunits in T conformation (continuous red line), fraction of subunits in ligand-bound R conformation (dashed black line), fraction of subunits in ligand-bound T conformation (dashed red line), fraction of ligand-free subunits in R conformation (dotted black line, highlighted with an arrow), and fraction of ligand-free subunits in T conformation (dotted red line). In addition, the fraction of ligand-bound subunits in either R or T conformation is shown (dashed green line). It is obvious that the contribution of subunits in T conformation to the ligand binding is very small (dashed red line). At very low ligand concentration, 77% of the protein subunits are in T conformation and 23% are in R conformation, according to the value of the equilibrium constant γ equal to 3.3. At low ligand concentration the total fraction of subunits in R conformation increases, while the total fraction of subunits in T conformation decreases due to the T → R conversion. However, both the fractions of ligand-bound subunits in R and T conformation increase due to ligand binding (although the increment in ligand-bound subunits in T conformation is negligible). At high ligand concentration the total fraction of subunits in R conformation further increases, while that of total subunits in T conformation decreases. And the fraction of ligand-bound subunits in R conformation further increases due to ligand binding, but the fraction of ligand-bound subunits in T conformation decreases due to the T → R conversion. Interestingly, the fraction of ligand-free subunits in R conformation dominates this region, with a maximal population of 60%. This is due to the concerted conversion of subunits within a given protein oligomer (all subunits within an oligomer undergo the conformational change T → R, even though not all of the subunits bind a ligand). At very high ligand concentration the fraction of ligand-bound subunits in R conformation dominates the conformational ensemble, reaching a maximal population of 100%. The R↔T equilibrium shows a switchover or crossover point (R and T are equally populated) at around 2 µM free bortezomib concentration. D) *Tt*ClpP peptidase activity was measured as a function of PKMamc concentration in the presence (15 µM) and absence of bortezomib.

Although the MWC model was employed forty years ago for the calorimetric study of the interaction of trout hemoglobin with carbon monoxide using a gas-liquid reaction calorimeter (*37*), the present work represents the first detailed mathematical description and implementation of the MWC model in ITC, for which the experimental methodology and the mathematical formalism are not the same as those briefly explained in that previous work.

The MWC model provides a rationale for *Tt*ClpP activation by bortezomib. The activation of *Tt*ClpP by the inhibitor bortezomib at moderate concentrations occurs as a consequence of the concerted conformational change within the protein tetradecamer, which makes the total fraction of subunits in the enzymatically active R conformation (F_R,Total_) much larger than the fraction of ligand-bound subunits in R conformation (F_R,Bound_); that is, there is no proportionality between F_R,Total_ and F_R,Bound_ (Fig. 5C). At intermediate bortezomib concentration, binding the fraction of inhibitor-free (active) subunits in R conformation is maximal and predominates in the speciation distribution (Fig. 5C). Thus, the extents of the cooperativity for binding and the cooperativity for activity are different (a phenomenon previously indicated for allosteric proteins in reference (*38*). If the conformational changes were not concerted, the activity of *Tt*ClpP would continuously decrease with increasing bortezomib concentration, with no maximal activity at intermediate bortezomib concentration. The increase in the fraction of inhibitor-free R subunits at moderate inhibitor concentration is the molecular event underlying the increase in activity. Similarly, the decrease in the fraction of inhibitor-free R subunits at high concentration of inhibitor is the molecular event underlying the decrease in activity at high inhibitor concentration.

Importantly, the ITC-derived model accurately predicts the activity data. According to peptide degradation activity measurements (Fig 1A), the maximal *Tt*ClpP activity is found at around 12.5 µM of total bortezomib. The ITC-derived MWC model predicts that the maximum fraction of inhibitor-free *Tt*ClpP in R conformation (F_R,Free_) is found at around 6 µM of free bortezomib (Fig 5C). Considering the population of bound *Tt*ClpP at 6 µM free bortezomib concentration (F_T or R,Bound_ =0.3; Fig. 5C), and taking into account the TtClpP subunit concentration of 10 µM (used in our activity measurements), one can calculate that the concentration of bortezomib bound to *Tt*ClpP is 3 µM. Consequently, the total concentration of bortezomib for achieving 6 µM free bortezomib (the concentration for maximal bortezomib-free subunits in R conformation) is around 9 µM, in excellent agreement with the activity measurements that reported maximum activity for 12.5 µM bortezomib concentration (Fig. 1A).

Furthermore, to test that the bortezomib induced R state has higher affinity for the substrate we measured peptide hydrolysis as a function of substrate concentration in the apo state and in the presence of non-saturating concentrations of bortezomib (15 µM). While there are differences between K_d_, a thermodynamic equilibrium constant, versus a purely kinetic constant as K_M_, in the presence of bortezomib *Tt*ClpP displayed an apparent K_M_ around 400 µM, and in its absence, no indications of substrate saturation was observed (Fig. 5D) reflecting the major differences between the *Tt*ClpP bortezomib bond and apo states. Together, the activity measurements and ITC data reveal that the activation of *Tt*ClpP by an inhibitor is achieved by shifting the conformational population towards a higher-affinity R state.

### Off-equilibrium MD simulations provide rationale for stabilization of extended state

The activity measurements, ITC data, X-ray structures and NMR spectroscopic data have established that by binding to a subset of *Tt*ClpP active sites, bortezomib as well as peptide-substrate favor a catalytically more active particle by keeping both the bound and free subunits in an active state, with the catalytic triad poised for catalysis. We attempted to bridge the functional and thermodynamic information from biochemistry and ITC with the structural view, by performing *in silico* experiments. Briefly, in these MD simulations we investigated the energetics of the transition from the extended (active) to the compressed (inactive) state, and interrogated whether the presence of a molecule bound to the active site alters the relative stability the two states. Near-equilibrium MD experiments were performed, in which the two heptameric rings were slowly “pushed” from the extended to the compressed state, using the centers of mass of the two rings as single variable for the steered-MD protocol (see Methods). Fig. 6A,B displays the force required for this extended-to-compressed transition, obtained from five independent in-silico experiments, as a function of the distance of the two half-rings, and Fig. 6B shows the work (i.e., the integrated force) for this process. These simulations have been performed either for the apo state (grey) or a sub-stoichiometrically peptide-loaded state. During the compression process none of the initially bound peptides were expulsed from their target catalytic sites and all the secondary structures were preserved (Fig. 6C-E). Interestingly, compressing ClpP from the extended conformation requires a stronger force in the presence of a sub-stoichiometric amount of alanine tripeptides, compared to the empty protein (Fig. 6A,B). These data provide a rationale for the enhanced stability of the extended conformation in the presence of ligand: even sub-stoichiometric amounts of ligand, *i.e.*, only partially occupied sites, lead to an overall stabilization of all subunits in the active state. The simulations were performed with peptides rather than bortezomib, due to the lack of reliable force field parametrization of boronic acid, and reveal the substrate-induced stabilization of the active extended state. It can be expected that this stability difference of extended over compressed state is even more pronounced with covalently bound bortezomib.

**Figure 6.**
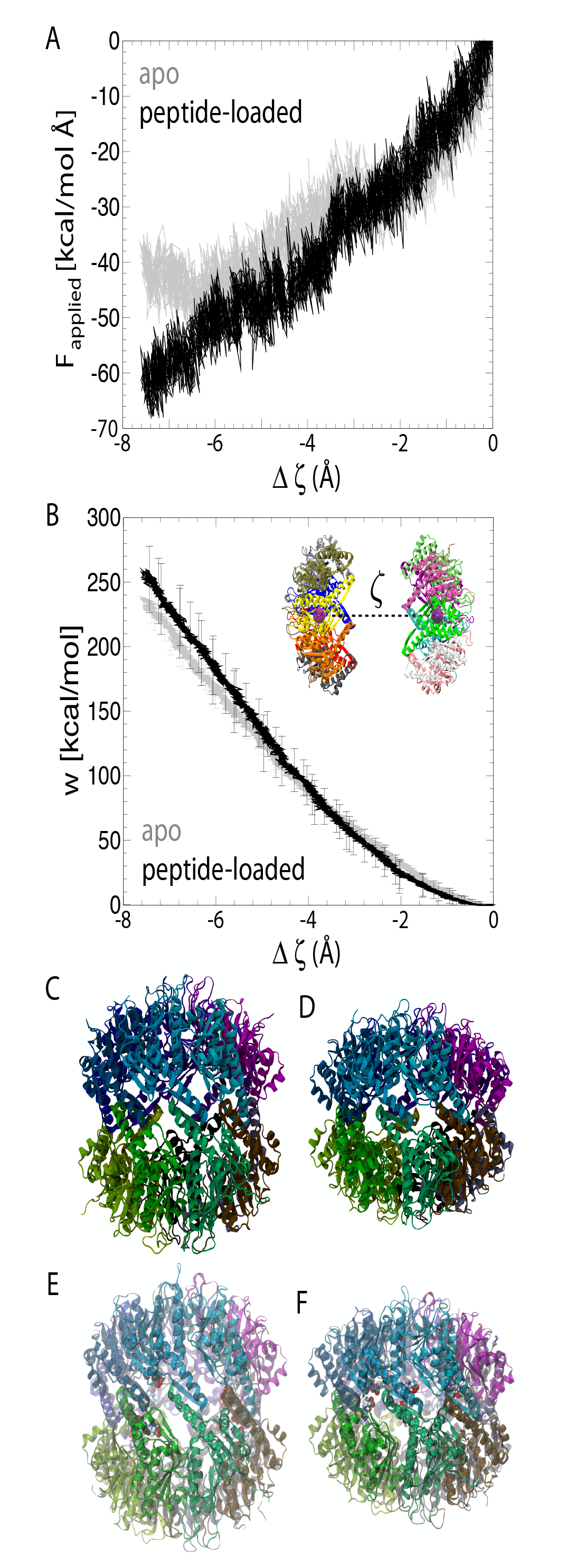
In silico pulling experiment of *Tt*ClpP from the extended to the compressed state. A,B) Profile of the force (A) and work (B) required for compressing the two rings towards each other as a function of the distance ζ between the rings (shown in the insert, with exaggerated distance for better visibility). C,D) Conformations of *Tt*ClpP extended (C) and *Sa*ClpP compressed (PDB: 3QWD) (D) states, observed in crystal structures. E) Starting conformation of *Tt*ClpP in the steered MD simulations (shown as sphere representation). F) Final conformation of the compressed state, at the conclusion of the steered MD. Note that the peptides are still bound and the structure is in good overall agreement with the compressed state from X-ray diffraction, shown in (D).

## Discussion

The activation of different ClpPs by previously identified inhibitors has been independently reported (*3, 39*), but so far no rationale for these paradoxical results had been proposed. Bortezomib, similarly to the previously described N-blocked peptide aldehyde *Mtb* activators, does not follow the canonical definition for an allosteric activator. That is, they bind to the protease active site(*12, 13*), and their effect is not universal to all ClpP homologues. For instance, we show here that although bortezomib activates *Tt*ClpP (up to 100 µM), it is in the same concentration range a good inhibitor of *Ec*ClpP. This is a rather puzzling result considering the high sequence and structural similarity between the two proteins (58 % sequence identity). We show here that ClpP activation by inhibitors derives from the intrinsically cooperative character of ClpP tetradecameric structure. Bortezomib, although blocking a number of active sites, can overall increase the catalytic rate by provoking a concerted transition in the complex that stabilizes the active state conformation. In line with this, the near-equilibrium MD simulations performed in our study suggest that compression of the extended-active conformation of *Tt*ClpP is hampered when a fraction of the catalytic sites are occupied by unspecific tripeptide substrates. In the presence of a stronger specific binder such as bortezomib, it could be reasonably anticipated that compression of the active conformation be even more hindered. A comparable mechanism is likely occurring for *Mtb*ClpP1P2 activation, since it has been shown that the peptide activator would sterically clash with a non-active ClpP1 complex, therefore forcing the complex into an active conformation (*39*).

The similarity between the bortezomib *Tt*ClpP structure and the structure of *Tt*ClpP with peptides in the active site strongly suggests that our findings reflect an inherent substrate-dependent activation mechanism of the enzyme. Through this mechanism, the presence of the substrate in the ClpP active site promotes the activation of the remaining subunits allowing efficient and processive protein degradation. Our results also support the previously described bidirectional cross-talk between chaperone binding and protease active sites (*12*, *17*, *21*, *39*, *40*), since bortezomib is no longer an activator in the presence of the co-chaperone *Tt*ClpX, and *Tt*ClpP partially saturated with bortezomib binds strongly to *Tt*ClpX. Significant differences were observed between the activation of *Tt*ClpP peptidase activity and proteolytic activity by bortezomib. Indeed, while peptidase activation displayed a bell-shaped curve, protease activation showed a linear rise in the range of the used bortezomib concentration. Although the process underlying bortezomib activation in the two activities should be common, important differences exist between the two assays. Protein degradation is strictly dependent on substrate access to the proteolytic chamber since bigger substrates cannot freely diffuse to the proteolytic chamber. Indeed, our results which show that bortezomib can activate protein degradation demonstrate, for the first time, that modulation of protein entry can also be achieved via binding to the ClpP active site, and not exclusively by ClpX or ADEP binding. Protein access to the proteolytic chamber leads to high local concentrations of substrate present at the ClpP active sites. Although bortezomib activates protein degradation at low concentrations, it must compete with high concentrations of substrate in the proteolytic activity assay, therefore explaining the requirement for higher concentrations of drug to achieve activation. The range of bortezomib concentrations used in our proteolytic activity assay likely corresponds to the rising part of the bell curve observed in the peptidase activity assay. This also explains why several ClpP inhibitors are 10-fold more potent in peptidase assays as compared to protease assays (*41*).

It is unclear why the action of bortezomib and activators is not uniformly conserved between different ClpPs. Differences in enzymatic activity are generally more difficult to decode than major structural differences because allosteric pathways often result from subtle changes in conformational dynamics that affect population distributions, which are not readily observed from comparisons of structures. For example, single mutations or truncations in *Ec*ClpP N-terminal have been shown to induce the reversible inactivation of ClpP (*33*) and using an indirect methodology based on the association of an *Ec*ClpP E14W mutant with *Ec*ClpA, the equilibrium constant between an *Ec*ClpP active state (R) and inactive state (T) was previously quantified as K_eq_=7.5 (37 °C)(*42*), a value similar to one we describe here for *Tt*ClpP K_eq_=3.3 (45 °C). This is particularly interesting as it suggests that in the absence of mutations *Ec*ClpP preferentially populates active states while *Tt*ClpP is preferentially inactive (*33*). The energy barrier between the two states is however rather small, with an energy gap of 0.7 kcal/mol between R and T conformations in *Tt*ClpP. Given the high sequence identity between *Ec*ClpP and *Tt*ClpP, the key to the difference in the equilibria between active and non-active states must result from small sequence changes. *Ec*ClpP, as well as *Sa*ClpP contain a conserved Q_131_XT/S_133_ motif at the tip of the turn between helix αE and the β-strand 6 in the handle domain (Fig. S7). Here, Gln131 makes hydrogen bonds with the η1 and η2 nitrogens of Arg170 on a neighboring subunit of the same heptamer ring (intra-ring), and with the backbone oxygen of Glu169 and ε1 oxygen of Gln123 on a monomer of the other ring (inter-ring). The side chain of the conserved Thr133 forms hydrogen bonds with the nitrogen atoms of Gln123 and Lys146 (inter-ring). Gln123 is part of the conserved HQP motif that includes the catalytic triad His122, while Arg170 is the vicinal residue of the catalytic triad Asp171. It is clear that this extended hydrogen-bond network restricts the degrees of freedom of the above-referred catalytic triad residues. Note that these residues are neither conserved in *Tt*ClpP nor in *Mtb*ClpP1. In *Tt*ClpP, Gln131 is replaced by a threonine (Thr131), which interacts with either the ε1 nitrogen of Arg170 (intra-ring) or with Gln123 (inter-ring) in the *Tt*ClpP:peptide or *Tt*ClpP:bortezomib structures respectively. Ser133 of *Tt*ClpP contacts Gln123 and Lys146 (inter-ring) in a similar way as *Ec*ClpP Thr133 (Fig. S7). The replacement of the conserved glutamine Q131 by the shorter threonine in *Tt*ClpP or serine in *Mtb*CpP1 leads therefore to modifications in the hydrogen network that normally sustains the extended helix αE. Structural and molecular dynamics studies in *Sa*ClpP have shown that these key residues are essential for keeping the long helix E in a straight conformation. In *Tt*ClpP the exclusive hydrogen bond acceptor character and shorter side chain of Thr131 does not allow the formation of the network that simultaneously coordinates two of the members of the catalytic triad. This reduced H-bonding capacity is expected to result in an increased flexibility in that region and likely to a shift in equilibrium to non-functional conformations, possibly picked up by the flexibility measurements reported here by MAS NMR and MD simulations (Fig. 5). We propose that bortezomib binding by preferentially stabilizing the active site (R form) shifts the entire population to a fully active enzyme.

Interestingly, a different hydrogen bond network is also observed for *Mtb*Clp1. In this case, Gln131 is replaced by a serine, while Thr133 is replaced by an alanine. When purified alone, ClpP1 is an inactive enzyme and the X-ray structure of the ClpP1 tetradecamer shows that the catalytic triad is not in the active state (*43*). However, in complex with ClpP2, ClpP1 has been shown to be fully active likely because *Mtb*ClpP2, which contains the conserved QFT motif at the tip of the helix αE, can stabilize ClpP1 by forming a similar network as observed in *Ec*ClpP(*3*). Cooperativity allows multimeric systems to rapidly adapt to changes in the environment by more efficiently converting a biochemical input – the substrate/ligand concentration – to a biochemical response. Cooperative response to incoming substrates has been reported for other machineries of the protein quality control system. Since protein degradation is key to cell homeostasis, several levels of regulation have evolved to prevent uncontrolled proteolysis that would be deleterious for the cell. Indeed, ClpP, as well as other complexes like the 20S proteasome, hide their active sites inside an inaccessible catalytic chamber in order to prevent uncontrolled proteolysis. This architecture is fine-tuned by ClpP specific co-chaperones, which add another layer of regulation in protease control, either by “opening” the axial pores of the cylinder and translocating unfolded substrates into the chamber and/or by directly activating the ClpP active sites(*2*). Several reports have shown that ClpX is able to revert the conformational inhibition of ClpP induced by chemical compounds (*12*) or even by mutations known to promote ClpP inactive states (*17*). The fact that one of these inactive-state promoting mutations, namely the Asp172Asn *Sa*ClpP mutant, corresponds to the native form of *Lm*ClpP1 and several other ClpPs suggests that this could be a relevant mechanism of *in vivo* ClpP regulation in these species (*44*). Our data clearly supports a similar mechanism of conformational control for *Tt*ClpP. Our observation that *Tt*ClpX as well as *Ec*ClpX (data not shown) can activate *Tt*ClpP suggests that this enzyme is conformationally attenuated, that is, in absence of the chaperone is in a latent inactive state. The non-physiological activation observed with bortezomib is both a consequence of the inherent cooperative behavior of the protease and the fact that in the absence of activators its predominant state is the non-active T state (Fig. 5C).

The small-molecule-induced conformational shift opens new possibilities for drug development, particularly as not only peptide-degradation but also protein degradation is stimulated, as we report here (Fig. 1C). In order for a molecule to achieve activation, a proper balance between binding, conformational change, and sufficiently low affinity to allow peptide or protein competition for the active site must be obtained.

In summary, using a host of biochemical, biophysical and structural approaches, we propose a mechanism for the activation of ClpP by inhibitors with a key element: a concerted activating conformational change in the supramolecular ClpP tetradecameric structure elicited by the binding of inhibitors at subsaturating concentrations.Hints to a similar allosteric behavior have also been reported for the proteasome in the presence of bortezomib (*44*): while bortezomib efficiently inhibits the chymotrypsin site of the proteasome, it was observed that it induced a paradoxical activation of the trypsin site of the complex (*45*).

## Materials and Methods

### Cloning and mutagenesis

*Tt*ClpP DNA was directly amplified from *Thermus thermophilus* HB8 DNA and cloned in to a pet41c (Novagen) expression vector with *Ned*I and *Xho*I restriction enzymes. The gene expressed contained at its C terminal the additional residues LEHHHHHHHH in order to allow affinity chromatography purification. Full length ClpX DNA was first cloned into a pet28 vector with *Ned*I and *Hind*III restriction enzymes with the additional MGSSHHHHHHSSGLVPRGSH residues at the N-terminal. While full length ClpX was insoluble, as occurred with other species homologues, removal of its N-terminal domain (1-54), allowed expression of soluble protein (∆*Tt*ClpX) (Glynn et al., 2009; Wojtyra et al., 2003). Removal of *Tt*ClpX N-terminal was done by Genecust using *Ned*I and *Hind*III as restriction enzymes. *Tt*ClpP and ∆*Tt*ClpX were expressed in *E. coli* Bl21RIL cells following ON expression at 20 °C with IPTG 1 mM. S97A mutation was inserted using QuickChange II site directed mutagenesis II from Thermo Fisher.

### Protein purification

GFPssrA was purified as previously described (*46*). Casein fluorescein isothiocyanate from bovine milk was obtained from Sigma (C0528) and was loaded into a G25 column (GE Healthcare) in order to remove free fluorescein as described (*46*). *Tt*ClpP and ∆*Tt*ClpX were purified using native NiNTA affinity chromatography followed by size exclusion chromatography in a 16/600 S200 200pg Superdex column. U-[^2^H,^15^N],Ile-δ1-[^13^CH_3_]*Tt*ClpP was expressed as described(*46*). When expressed in D_2_O *Tt*ClpP was insoluble and the protein was recovered from *E. coli* inclusion bodies using denaturing conditions. Briefly, cells were harvested and resuspended in denaturing buffer containing 8 M urea followed by sonication and centrifugation to remove cell debris. The resulting supernatant was loaded into a NiNTA column and washed several times with denaturing buffer. The column was then equilibrated in refolding buffer (NaPi 100 mM pH 8, glycerol 5%, 100 mM NaCl) followed by elution with elution buffer (NaPi 100 mM pH 8, glycerol 5%, 100 mM NaCl and 400 mM imidazole). The eluted fractions were loaded into a 16/600 S200 200pg Superdex column and the fractions corresponding to native ClpP were pulled. *Ec*ClpP was purified as described in(*47*). Ftsz was a kind gift of Dr Tomas Hosek and was purified as previously described(*47*).

### Biochemical Assays

PKMamc degradation by *Tt*ClpP and *Ec*ClpP was measured as described(*41, 48*). Briefly *Tt*ClpP in Hepes pH 7.6 50 mM NaCl 50 mM was mixed with indicated concentrations of PKMamc and bortezmib/Bz-LL (or respective DMSO volume) at 60 °C (or indicated temperature) and fluorescence (340 nm excitation, 462 emission) was measured at defined time intervals. FITC casein degradation by *Tt*ClpP was measured as described(Akopian et al., 2012); in a typical assay FITC-casein (0.15 µM) was mixed with *Tt*ClpP_14_ (0.1 μM) and fluorescence (440 nm excitation, 509 nm emission, cut off 495 nm) increase was monitored. GFPssrA (5.5 μM) degradation by *Tt*ClpX_6_ (0.2 μM) and *Tt*ClpP_14_ (0.1 μM) in the presence of ATP (10 mM) was monitored taking advantage of GFPssra intrinsic fluorescence (440 nm excitation, 509 nm emission, cut off 495 nm) at the indicated temperatures. FtsZ degradation by *Tt*Clp*P* was measured at 37 °C. Briefly *Tt*ClpP_14_ (1 μM) was mixed with purified FtsZ (20 μM) in the presence of the indicated concentrations of bortezomib. Aliquots were removed at the indicated time points and the samples were loaded in a 15% SDS agarose gel. *Tt*ClpP reaction with ActivX TAMRA-FP was executed according to manufacturer instructions (Thermo Fisher Scientific). *Tt*ClpP_14_ (1 mg/ml) was incubated in the indicated conditions with 1 μl of TAMRA-FR reagent (final concentration 5 μM) with the indicated conditions. At defined time intervals aliquots were removed and loaded in a 4-20% SDS gel. Gels were visualized in a Biorad ChemiDoc XRS system. *Tt*ClpP (20 μM) unfolding was measured in a Varian Cary Eclipse Spectrofluorimeter monitoring tryptophan intrinsic fluorescence (280 nm excitation, 350 nm emission). *Tt*ClpP_14_ analytical SEC was done using a Superdex 75 10/300 GL.

### NMR experiments

Magic-angle spinning solid-state NMR experiments were performed on a Bruker Avance III spectrometer with a 1.3 mm HCND MAS NMR probe, at an MAS frequency of 55 kHz and an effective sample temperature of 30 °C. Backbone resonance assignment was achieved with proton-detected triple-resonance experiments described earlier(Fraga et al., 2017). The assignment was obtained for 97 residues; many more spin systems were observable in the spectra, but spectral overlap challenged further assignment, which likely requires additional high-dimensional experiments. Given that in the present context we were interested in probing interactions and dynamics, the current extent of assignment was sufficient here.

For the measurement of chemical-shift perturbation upon addition of bortezomib, we prepared two samples of u-[^2^H,^13^C,^15^N]-labelled ClpP in H_2_O-based buffer (100 mM Tris pH 8.5, glycerol 5%, NaCl 100 mM) containing either 10 mM mM bortezomib and 5 % DMSO, or only 5 % DMSO. In both cases, the protein (10 mg/mL) was sedimented into a 1.3 mm Bruker MAS NMR rotor, and hCANH spectra were collected in ca. 2.5 days per spectrum. The peak positions in the two spectra were extracted for all assigned backbone sites using CCPN (*49*).

Spin-relaxation measurements (^15^N R_1ρ_) were performed with proton-detected 3D hCANH experiments. The 3D approach was chosen in order to resolve sites which would be overlapped in 2D HN spectra. The experiment comprised a ^15^N spin-lock element of 15 kHz radiofrequency field strength and delays of 2 ms and 55 ms. Total experimental times were 3 days per 3D experiment. All spectra were with Topspin v3.5 and analyzed with CCPN. Relaxation rate constants were obtained using in-house written Python scripts using the peak intensities extracted from the 3D spectra, and error bars were determined from Monte Carlo analysis based on the spectral noise level (determined in CCPN).

Solution-state NMR experiments were performed on a Bruker Avance III spectrometer, equipped with cryogenically cooled TCI probeheads, operating at a magnetic field strength corresponding to ^1^H Larmor frequencies of 700 MHz. The sample temperature was set to 60 °C and two-dimensional (2D) SOFAST-methyl-TROSY NMR experiments were recorded with an adjusted duration depending on the final concentration of the proteins (experimental time ranging from 10 to 120 min per sample)(*50*). The angle of the proton excitation pulse was set to 30°, and the recycling delay was optimized to 0.6 s to achieve the highest sensitivity (*49*). 250 μM of U-[^2^H,^15^N], Ile-δ1-[^13^CH_3_]*Tt*ClpP and U-[^2^H,^15^N],Ile-δ1-[^13^CH_3_]S97A*Tt*ClpP were mixed with 2 mM bortezomib (or DMSO equivalent) in Tris buffer (pH 8.5 mM 100 mM, NaCl 100 mM and glycerol 5%).

### Molecular Dynamics simulations

#### Computational assays

The three-dimensional structure utilized in the simulations was obtained from apo-TtClpP, which corresponds to an extended conformation of the protein, bereft of substrate. The missing N-terminal β-hairpin residues were added, employing Modeler 9.17. Positioning of the seven loops was done concomitantly for each of the upper and the lower hemispheres of the protein. For the protein with bound peptide substrates, use was made of the apo-TtClpP structure featuring stretches of electron density characteristic of short peptides, over which zwitterionic alanine tripeptides were superimposed. The computational assays consisted of the extended *Tt*ClpP, with or devoid of alanine tripeptides bound to the 14 serine active sites of the protein, immersed in a bath of 75,307 water molecules, with a 0.16 M NaCl concentration, corresponding to a cubic simulation cell of length equal to 138 Å at thermodynamic equilibrium.

#### Molecular dynamics protocol

All the MD simulations reported in this study were performed using NAMD 2.12, with the CHARMM36 force field for proteins and lipids, and the TIP3P water model. A Langevin thermostat with a damping coefficient of 1 ps^−1^ maintained the temperature at 27°C. The Langevin piston method was employed to keep the computational assay at a nominal pressure of 1 atm. Covalent bonds involving hydrogen atoms were constrained to their equilibrium length by the Rattle algorithm. The Settle algorithm was utilized to constrain water molecules to their equilibrium geometry. Long-range electrostatic forces were evaluated using the particle-mesh Ewald algorithm with a grid spacing of 1.2 Å, while a smoothed 12-Å spherical cutoff was applied to truncate short-range van der Waals and electrostatic interactions. The r-RESPA multiple time-stepping algorithm combined with a mass-repartitioning scheme was employed to integrate the equations of motion with an effective time step of 4 fs for short-range interactions and of 8 fs for long-range electrostatic interactions.

### Simulation design

The extended apo-TtClpP was probed in a 1-µs MD simulation. In addition, the computational assay consisting of the protein with zwitterionic alanine tripeptides bound to its 14 serine active sites was examined in three independent, 1-µs MD simulations, using distinct initial conditions. These simulations are aimed at exploring how the presence of the substrate impacts the flexibility of *Tt*ClpP, alongside the relevance of the alanine tripeptide as a proxy for the peptide chain that binds to the Ser97 active site.

### Near-equilibrium pulling experiments

To address how the occupancy of the serine active sites of *Tt*ClpP affects the compressibility of *Tt*ClpP, we have examined the conformational transition of the protein from its extended state to its compressed state by means of near-equilibrium pulling experiments. Toward this end, a steered MD protocol was employed, pulling at constant velocity a stiff spring of force constant equal to 100 kcal/mol Å. The transition coordinate was defined as the Euclidian distance separating the center of mass of the upper hemisphere to that of the lower hemisphere, computed from the position of all α-carbon atoms. The length of the transition coordinate was determined based on apo-*Tt*ClpP and the structure of the compressed conformation (PDB: 3QWD), and amounts to 8 Å. Three independent, 200-ns long realizations were performed for the empty protein and that containing the zwitterionic alanine tripeptides, pulling irreversibly the two lobes of the protein from an extended state to a compressed one.

### Isothermal Titration calorimetry

The interaction between *Tt*ClpP and bortezomib was assessed by ITC using an Auto-iTC200 (MicroCal-Malvern Instruments, Malvern UK). Calorimetric titrations were performed with a 1.4 mM bortezomib solution in the injecting syringe and a 10 µM ClpP solution in the calorimetric cell in Hepes pH 7.6 50 mM, NaCl 50 mM. All solutions were properly degassed to avoid bubble formation during stirring. In each titration, a sequence of 19 2-µL injections was programmed, with reference power of 10 µcal/s, initial delay of 60 s, spacing between injections of 150 s, and stirring speed of 750 r.p.m.

The heat evolved after each ligand injection was obtained from the integral of the calorimetric signal. Experiments were performed in replicates and data were analyzed using in-house developed software implemented in Origin 7 (OriginLab, Northampton, MA). The binding isotherms (ligand-normalized heat as a function the molar ratio) were analyzed considering the Monod-Wyman-Changeux (MWC) model for ClpP, an oligomeric macromolecule consisting of 14 identical subunits containing a single ligand binding site each (*50*). Non-linear least squares regression allows determining the binding parameters: equilibrium constants (K_R_, K_T_, γ), enthalpies (ΔH_R_, ΔH_T_, ΔHγ), and fraction of active protein (N). A fitting routine was implemented in Origin 7.0 (OriginLab).

### MWC model implemented in isothermal titration calorimetry

The Monod-Wyman-Changeux (MWC) model considers an oligomeric macromolecule consisting of n identical subunits containing a single ligand binding site each. Those subunits may adopt two conformations, R (relaxed) and T (tense) with different ligand binding affinities (K_R_ > K_T_). Within a given oligomer, all subunits exhibit the same conformation and ligand binding occurs to any subunit independently. However, although T state predominates initially, as a consequence of the higher ligand binding affinity for R conformation, ligand binding will shift the conformational equilibrium by indirectly eliciting a concerted conformational change of all subunits towards R conformation within the same oligomer at once, resulting in a cooperative binding behavior. The MWC model can only reproduce positive cooperativity, while the Koshland-Nemethy-Filmer (KNF) model can reproduce both negative and positive cooperativity. Both models, MWC and KNF, represents reduced and limited versions of a general allosteric model (*38, 51*) that takes into account all possible conformational states (e.g. mixed conformations within a protein oligomer) and all possible liganded states, but, in practice, they are considerably more useful and manageable than the general allosteric model.

The binding polynomial for a macromolecule with n subunits that can exist, all at once within a given oligomer, in two different conformations, R and T, and each subunit has a single ligand binding site is given by:

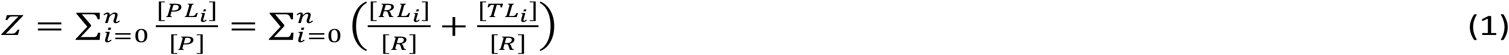

where PL_i_ represents the protein complex with i ligand molecules bound, and RL_i_ and TL_i_ refer to the complexes of each oligomeric conformational state with i binding sites occupied by ligand molecules. In terms of site-specific binding parameters, the MWC binding polynomial is written as follows:

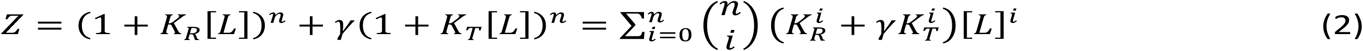

where K_R_ and K_T_ are the site-specific microscopic intrinsic association constants for a binding site in a protein subunit in the R and T state, respectively, with K_R_ > K_T_, and γ is the equilibrium constant for the conformational equilibrium between the T and R oligomers (γ = [T_n_]/[R_n_]). Because initially the oligomer in T conformation predominates, the equilibrium constant γ is larger than 1. Each term represents the binding sub-polynomial considering n independent ligand binding sites in each conformational state, R and T.

Valuable information can be extracted from the binding polynomial. In particular, the molar fraction of each liganded species:

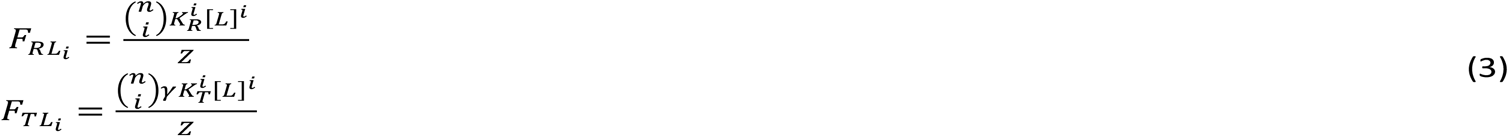

The two first derivatives of the binding polynomial respect to ligand concentration and temperature provide two fundamental quantities, the average number of ligand molecules bound per oligomeric macromolecule:

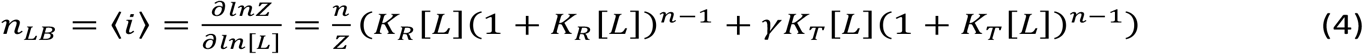

that can be expressed more conveniently as:

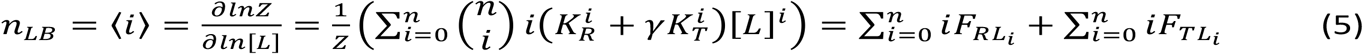

and the average molar excess binding enthalpy:

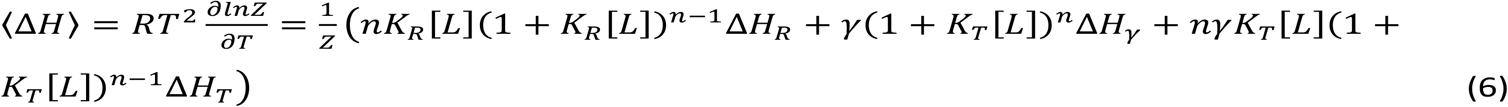

that can also be expressed more conveniently as:

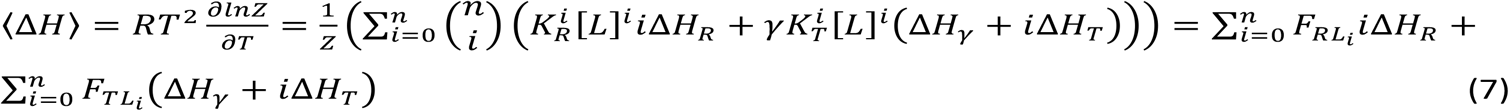

where ΔH_R_ and ΔH_T_ are the site-specific microscopic intrinsic ligand binding enthalpies for a binding site in a protein subunit in the R and T state, respectively, and ΔHγ is the enthalpy associated to the concerted conformational change between R and T states.

The binding equations corresponding to the binding equilibrium derive from the mass conservation and the chemical equilibrium:

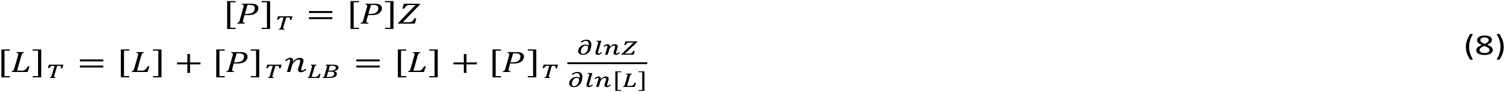

The last equation can be transformed into a (n+1)th-degree polynomial equation in [L] (in this case, a 15th-degree polynomial equation) with coefficients that are functions of K_R_, K_T_, γ, [P]_T_, and [L]_T_, and that can be solved numerically (e.g. Newton-Raphson method) for the unknown [L]. Eq. 8 must be solved for each experimental point in the calorimetric titration (i.e., each ligand injection j), for which the total concentrations of protein and ligand after each injection j are calculated as follows:

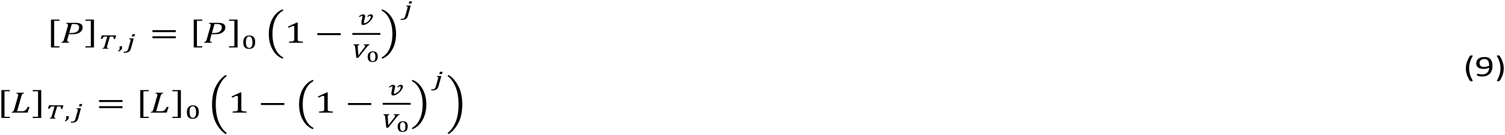

where [P]_0_ is the initial macromolecule concentration in the cell, [L]_0_ is the ligand concentration in the syringe, V_0_ is the calorimetric cell volume, and v is the injection volume. Once the free ligand concentration is known, the concentration of each complex after each injection can be calculated (subscript j omitted for the sake of clarity):

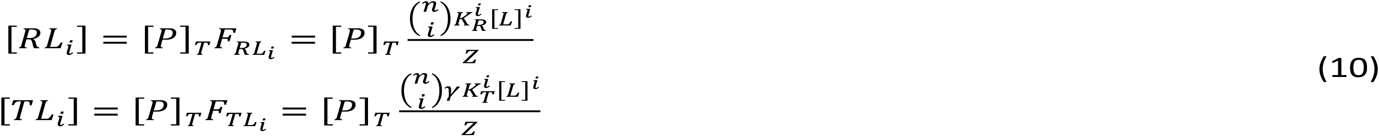

The heat effect, q_j_, associated with each injection j is calculated considering it reflects the change in the average excess molar binding enthalpy or in the concentration of all complexes in the calorimetric cell between injection j and j-1:

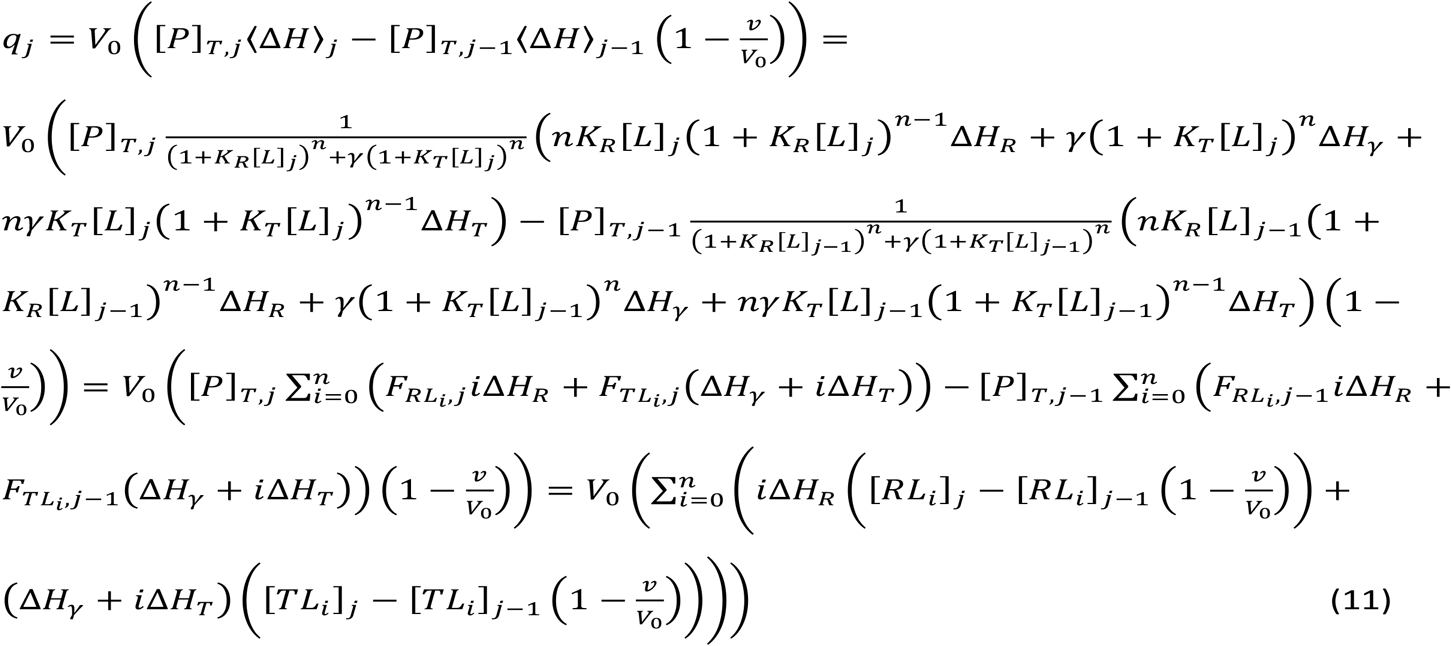

Finally, q_j_ is normalized by the amount of ligand injected during each injection, and an adjustable parameter q_d_ accounting for the background injection heat (due to solution mismatch, turbulence, etc.) is also included:

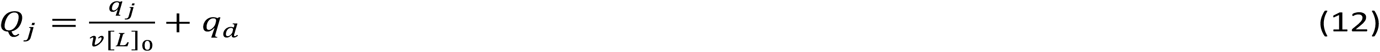

Additionally, a normalizing parameter N is included in eq. 9 multiplying [P]_0_ in order to account for the active or binding-competent fraction of macromolecule (percentage of protein able to bind ligand).

Non-linear least squares regression allows determining the binding parameters K_R_, K_T_, γ, ΔH_R_, ΔH_T_, ΔHγ, N and q_d_. A fitting routine was implemented in Origin 7.0 (OriginLab).

For the sake of simplicity, a non-normalized binding polynomial has been employed in eq. 2, where only the ligand-free R state has been taken as the reference state. A renormalized binding polynomial can be constructed taking the subensemble of ligand-free states, R and T, as the reference (ensemble) state, resulting in a standard binding polynomial with a leading term equal to 1 when grouping terms according to powers in [L]. The new binding polynomial is given by:

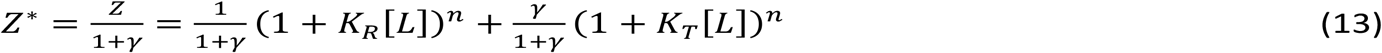

and the only difference in the subsequent development is that a different reference value for the excess average ligand binding enthalpy is employed:

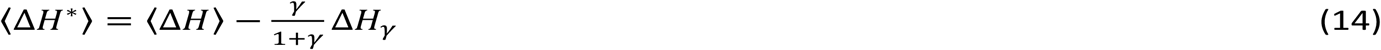

The second term in the right-hand side of eq. 14 after the minus sign is a constant value equal to the average excess enthalpy of the non-liganded fraction of protein (that is, ligand-free R and T states):

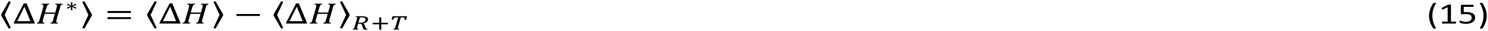

Thus, that term introduces a shift in the enthalpy scale, and an additional constant term in the average excess molar enthalpy makes no difference when calculating the heat effect for each injection, as can be easily proven:

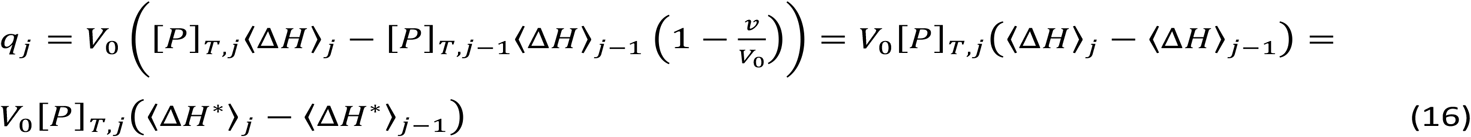

A nested MWC cooperativity model for the behavior of ClpP in which each heptamer might undergo a specific concerted conformational change^62^ could have been applied to ClpP, but that would have added three additional equilibrium constants and three additional enthalpy changes, which would result in overparameterization in the fitting function and correlation/dependency between fitting parameters. Also, the KNF cooperativity model in which sequential conformational changes are occurring as the oligomer is being occupied by the ligand^62^ could have been applied to ClpP, but, again, more parameters should have been considered in the fitting function, resulting in overparameterization. Besides, the KNF model cannot reproduce the activation effect induced by bortezomib, because in the KNF model the conformational change induced by the ligand is specifically restricted to those subunits binding the ligand, with no possibility of ligand-free subunits undergoing the activating conformational change. Thus, the MWC cooperativity model is the minimal model able to reproduce the behavior of ClpP.

The concentration (and the evolution along a titration) of the different protein species (R and T conformations) can be readily calculated from the previous equations. The total fraction of protein subunits in the conformation R or T, F_R,T_ and F_T,T_, respectively, are given by:

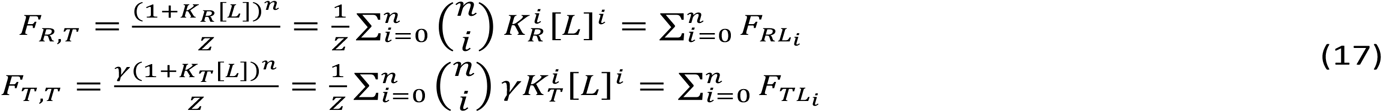

and the fraction of protein subunits in the conformation R or T that are bound to the ligand, F_R,B_ and F_T,B_, respectively, are given by:

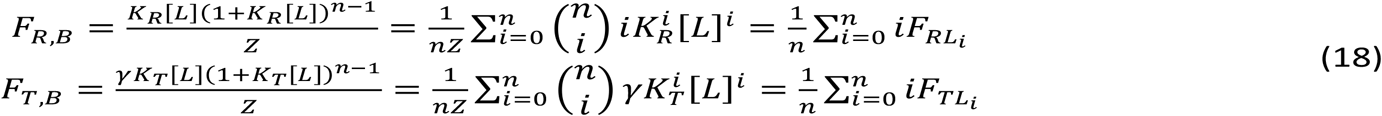

The fraction of ligand-free protein subunits in the conformation R or T, F_R,F_ and F_T,F_, can be calculated as the following differences:

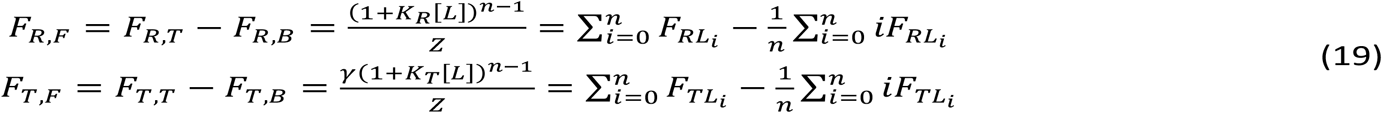

### Crystallization of *Tt*ClpP and the *Tt*ClpP:bortezomib complex

Crystallization trials were set up manually in a sitting drop vapour diffusion set-up using 24-well sitting drop plates (Hampton research) with drop volumes of 1 µl (0.5 µl protein solution + 0.5 µl reservoir solution). Purified *T. thermophilus* ClpP was concentrated to 9 mg/ml. Approximately one week after setting up crystal plates at 293 K, large cube shaped crystals were obtained in a condition containing 0.1 M Sodium acetate pH 4.8 and 30% PEG 400.

To obtain crystals of the *Tt*ClpP:bortezomib complex, prior to setting up crystal plates, 5 µl of a 100 mM bortezomib solution dissolved in DMSO was added to 45 µl of *Tt*ClpP solution (9 mg/ml) resulting in a final concentration of 10 mM bortezomib and 8.1 mg/ml of *Tt*ClpP. Large brick shaped crystals of the *Tt*ClpP:bortezomib complex were obtained at 293 K in the same condition as *Tt*ClpP (0.1 M Sodium acetate pH 4.8, 30% PEG 400). *Tt*ClpP crystals were scooped, transferred to a solution containing 32 % PEG 400 and 15 % glycerol, and subsequently flash-cooled in liquid nitrogen. Crystals of the *Tt*ClpP:bortezomib complex were scooped directly from the drops and flash-cooled in liquid nitrogen. Diffraction data of *Tt*ClpP (1.95 Å) and the *Tt*ClpP:bortezomib complex (2.70 Å) were collected at the ID30A – MASSIF (*Tt*ClpP) and ID23-1 MX (*Tt*ClpP:bortezomib) beamlines at ESRF, Grenoble, France. The obtained datasets were processed using the XDS package (*52*).

### Structure determination of *Thermus thermophilus* ClpP and the *Tt*ClpP:bortezomib complex

Diffraction data for *Tt*ClpP was processed in spacegroup C2 (a = 105.98 Å, b = 162.79 Å, c = 107.95 Å, α = γ = 90 °, β = 116.34 °). The structure was solved by maximum-likelihood molecular replacement (MR) using Phaser (*53*) in Phenix (*54*). The X-ray structure of *Ec*ClpP in complex with ADEP1 was used as a search model (PDB id: 3KTI(*55*)). Before running Phaser, side-chains of the search model were trimmed using the Schwarzenbacher (Schwarzenbacher et al., 2004) sidechain pruning method in Sculptor (Phenix). The resulting MR solution was refined in Phenix using positional (XYZ) and Real-space refinement with non-crystallographic symmetry (NCS) restraints, occupancy refinement, individual ADP refinement with TLS and optimized X-Ray/stereochemistry and X-Ray/ADP weights. Multiple rounds of refinement in Phenix were followed by additional manual model building carried out in Coot (*56*).

Diffraction data for *Tt*ClpP:bortezomib was processed in spacegroup C222_1_ (a = 135.14 Å, b = 168.74 Å, c = 166.08 Å, α = β = γ = 90 °). The structure was solved by maximum-likelihood molecular replacement (MR) using Phaser in Phenix, taking the refined 1.95 Å *Tt*ClpP structure as a search model. Refinement of the *Tt*ClpP:bortezomib structure in Phenix was carried out using the same strategy as for *Tt*ClpP (positional and real-space refinement with NCS restraints, occupancy refinement, individual ADP refinement with TLS and optimized X-Ray/stereochemistry and X-Ray/ADP weights, followed by additional manual model building in Coot).

### Data Availability

Coordinates and structure factors for *Thermus thermophilus* ClpP in complex with peptide or bortezomib have been deposited in the Protein Data Bank (PDB) with accession codes 6HWM and 6HWN respectively.

## Acknowledgments

This work used the platforms of the Grenoble Instruct center (ISBG; UMS 3518 CNRS-CEA-UJF-EMBL) with support from INSTRUCT (“Innovative EM/NMR approach for the characterization of the drug target ClpP APPID: 301“), FRISBI (ANR-10-INSB-05-02) and GRAL (ANR-10-LABX-49-01) within the Grenoble Partnership for Structural Biology (PSB). Special thanks to Dr Tomas Hosek (Institute Biologie Structurale) for FtsZ and Dr Tatos Akopian (Harvard School of Public Health, USA), for testing peptide substrates. We thank the ESRF for beamtime at ID30A and ID23A. This work was supported by Spanish Ministerio de Economia y Competitividad (BFU2016-78232-P), and Instituto de Salud Carlos III co-funded by European Union (PI15/00663, ERDF/ESF, “Investing in your future”) (PI15/00663).

---

**Figure Supplement 1.**
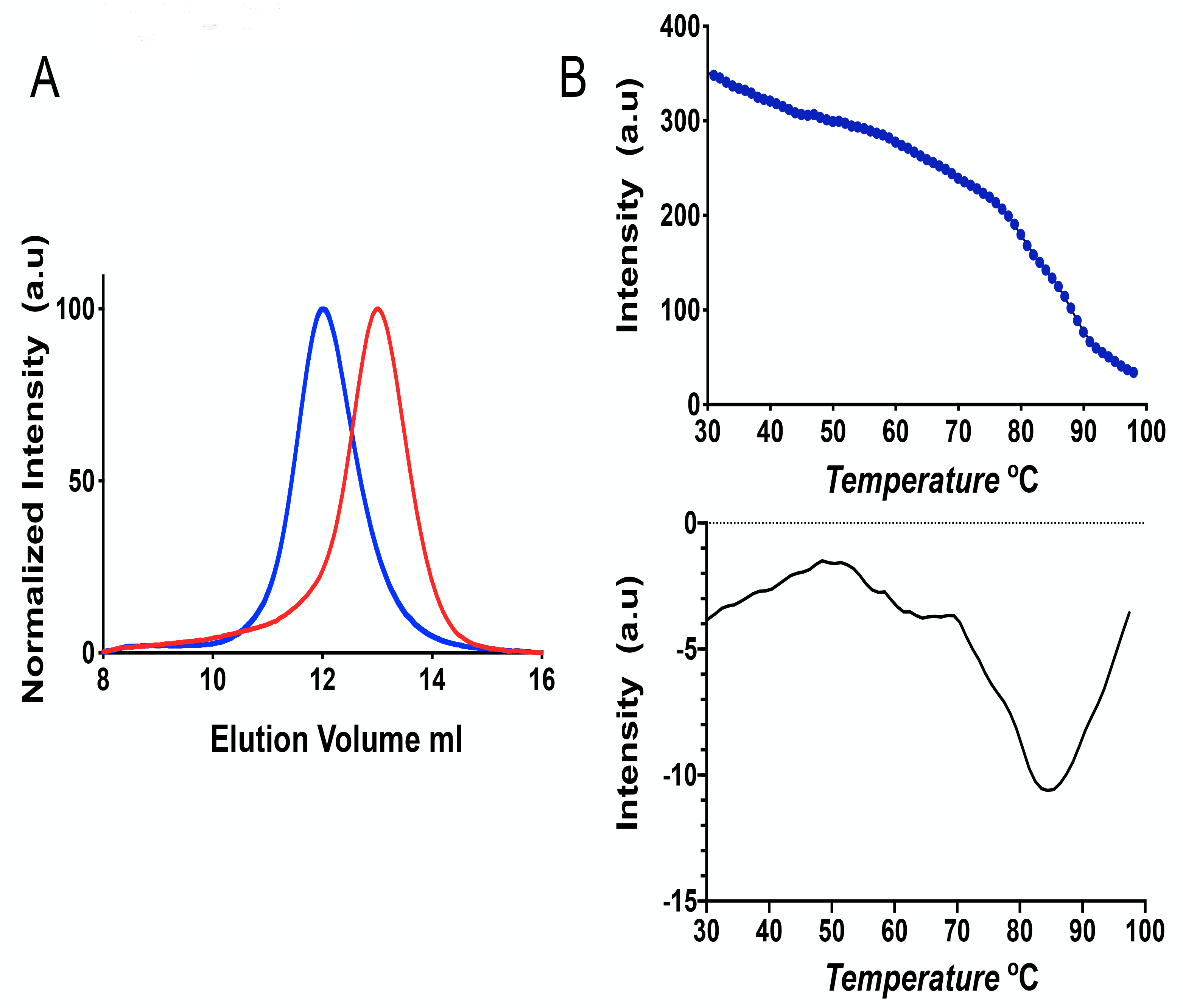
A) *Tt*ClpP eluted as a dodecamer in a low ionic strength buffer (100 mM NaCl) in a Superdex 75 10/300 GL (blue curve). In the presence of 300 mM (NH_4_)_2_SO_4_ smaller species were observed consistent with previously reported heptameter complexes (red curve). B) The indole group of tryptophan is very sensitive to its environment. Plot of *Tt*ClpP intrinsic fluorescence as a function of temperature (upper panel). The melting point of TtClpP (84°C) was estimated using the first derivative of the denaturation curve (lower panel).

**Figure Supplement 2.**
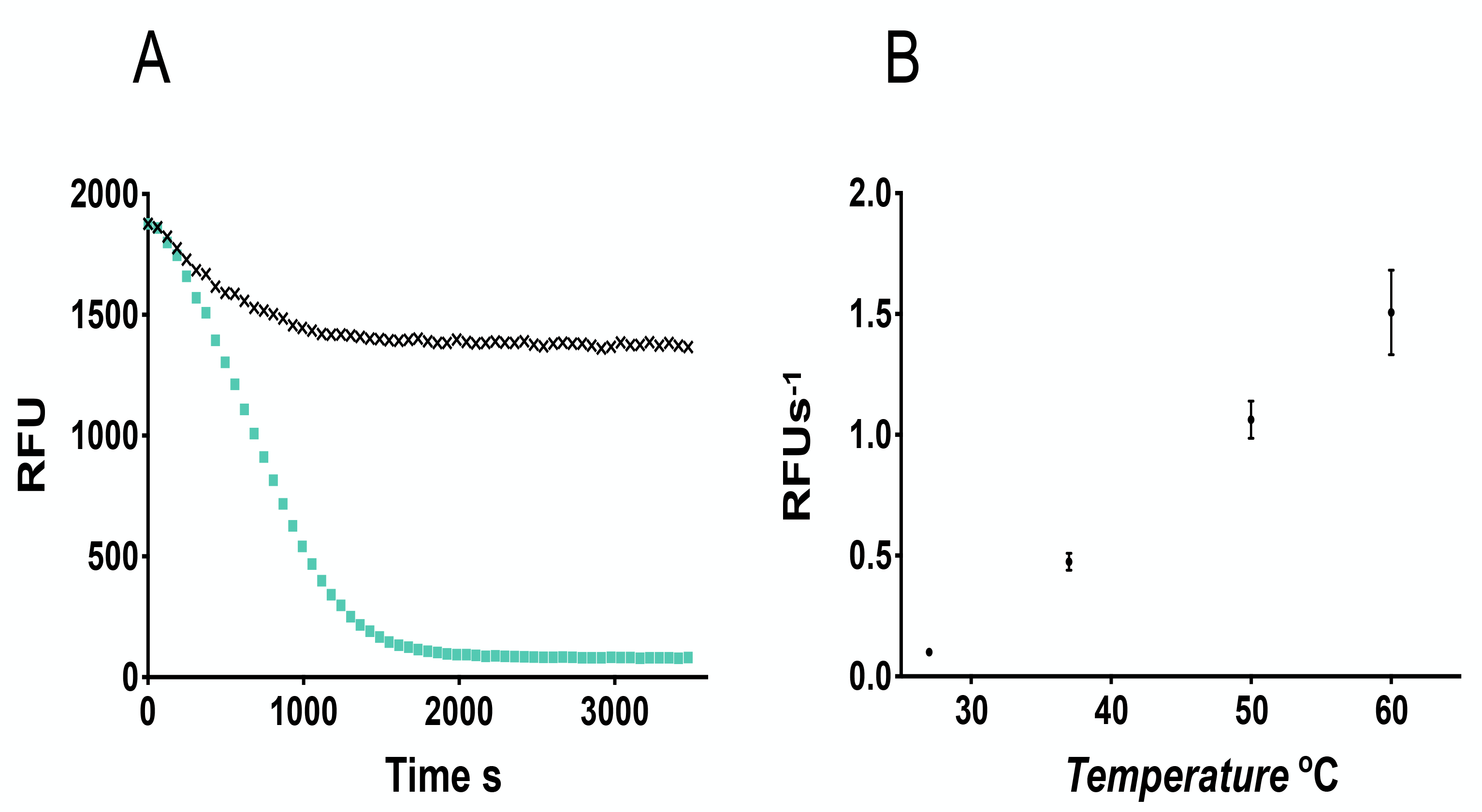
A) Stable proteins like GFPrrA cannot by degraded by *Tt*ClpP alone as they cannot access the proteolytic chamber. *Tt*ClpP can, in the presence of *Tt*ClpX and ATP, catalyse the degradation of GFPssra (cyan squares). In the absence of ATP no degradation is observed (black squares). The small drop in fluorescence observed in the absence of ATP results from temperature equilibrium (assay at 60 ºC). B) Initial rates for GFPssrA degradation by *Tt*ClpXP were plotted as a function of the temperature used. Maximum activity was observed at 60 ºC, the maximum incubation temperature allowed in the available fluorimeter.

**Figure Supplement 3.**
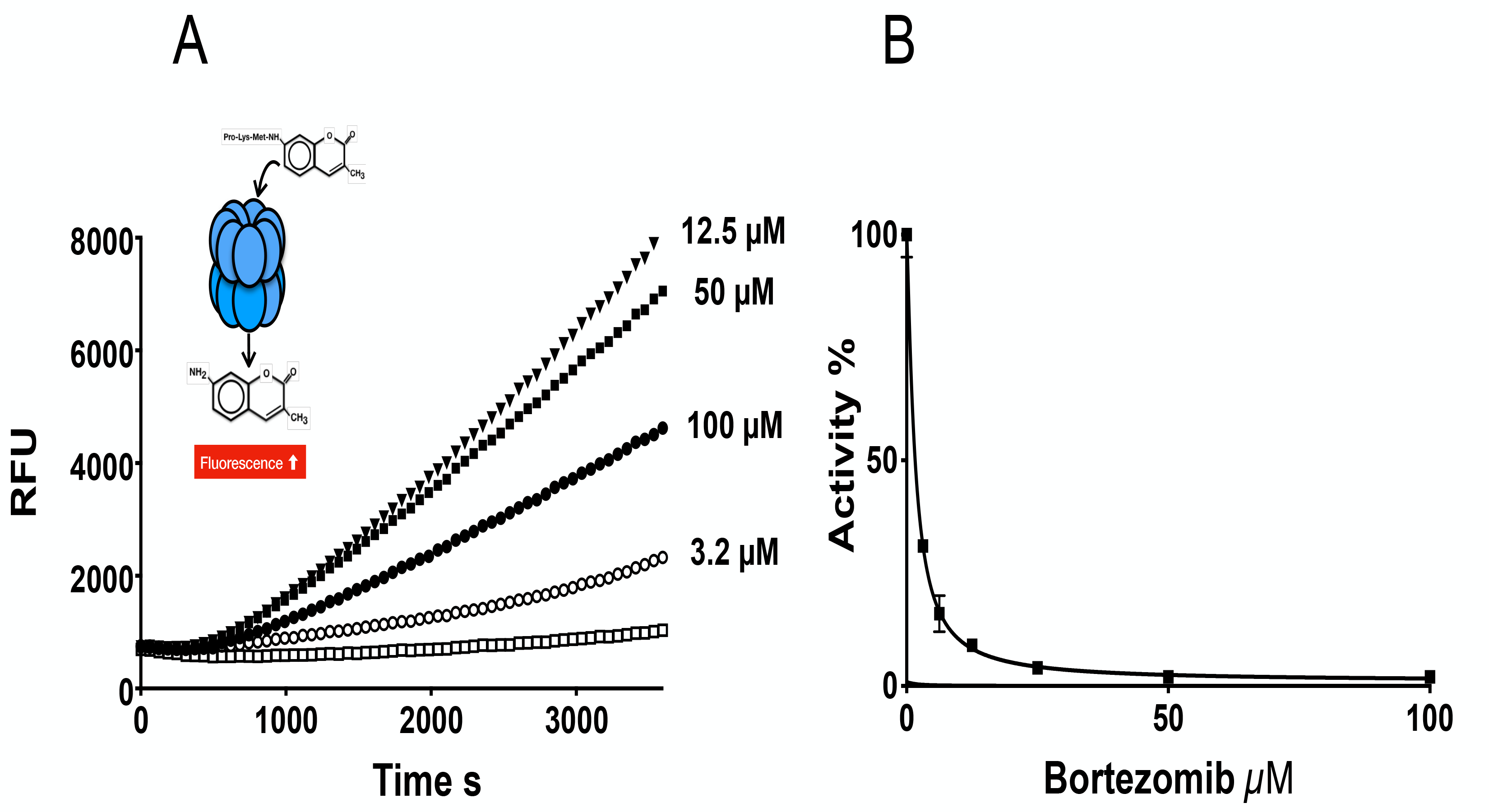
A)*Tt*ClpP peptidase activity was measured with the substrate PKM-amc (100 µM) in the presence of bortezomib at the indicated concentrations. As the peptide is cleaved, 7-Amino-4-methylcoumarin is released resulting in an increase in the fluorescence measured. B) Bortezomib is an inhibitor of *Ec*ClpP peptidase activity with an IC_50_ of 1.6 ± 0.1 µM.

**Figure Supplement 4.**
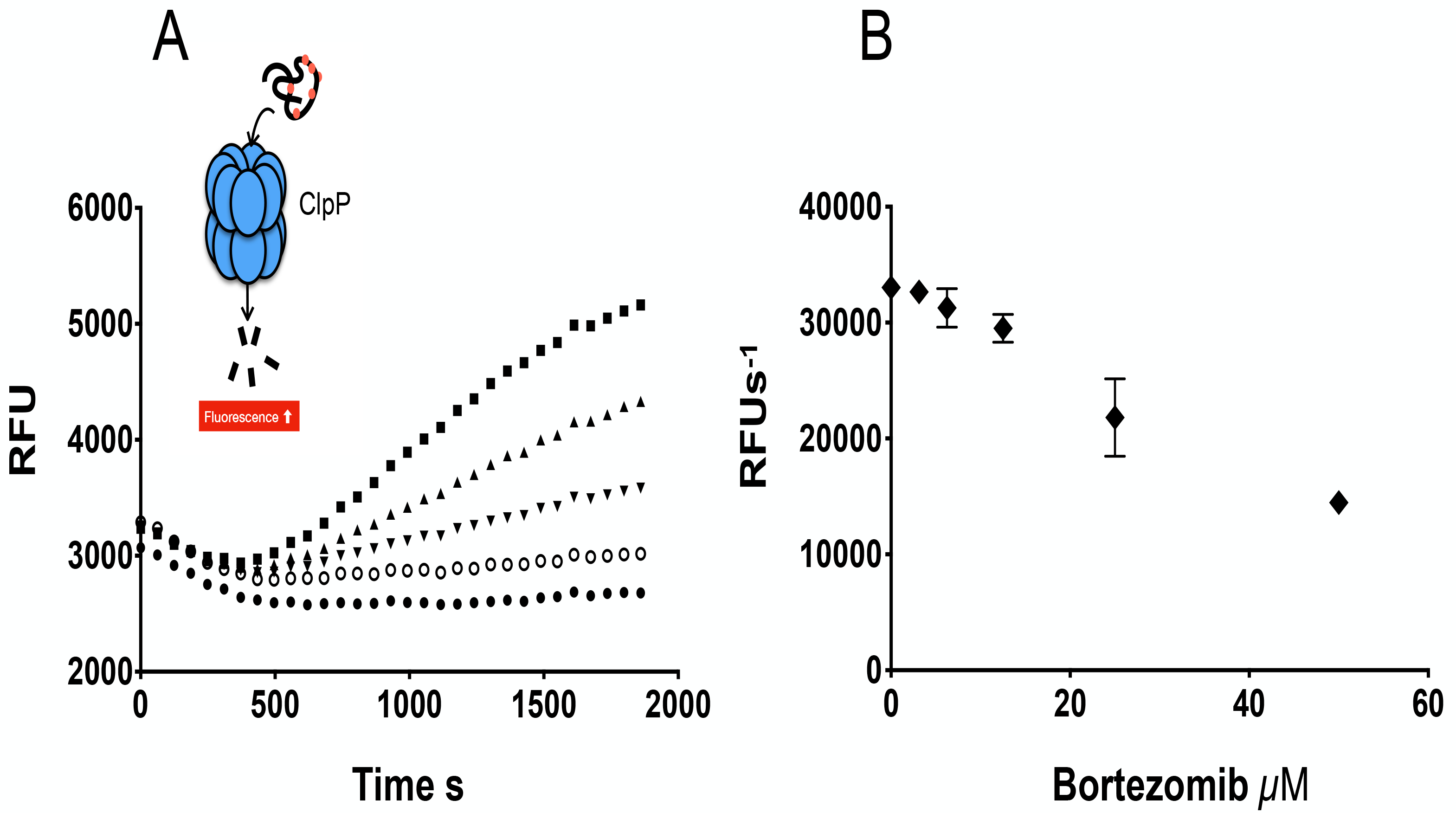
A) The degradation of the unfolded protein substrate FITC-casein by TtClpP was measured in the presence of the 100 μM square), 50 µM (triangle), 25 µM (inverted triangle), 6.25 µM (empty circle) and control (filled circled). B) In the presence of TtClpX (equimolar), bortezomib does not activate TtClpP and instead an inhibition of the peptidase activity is observed.

**Figure Supplement 5.**
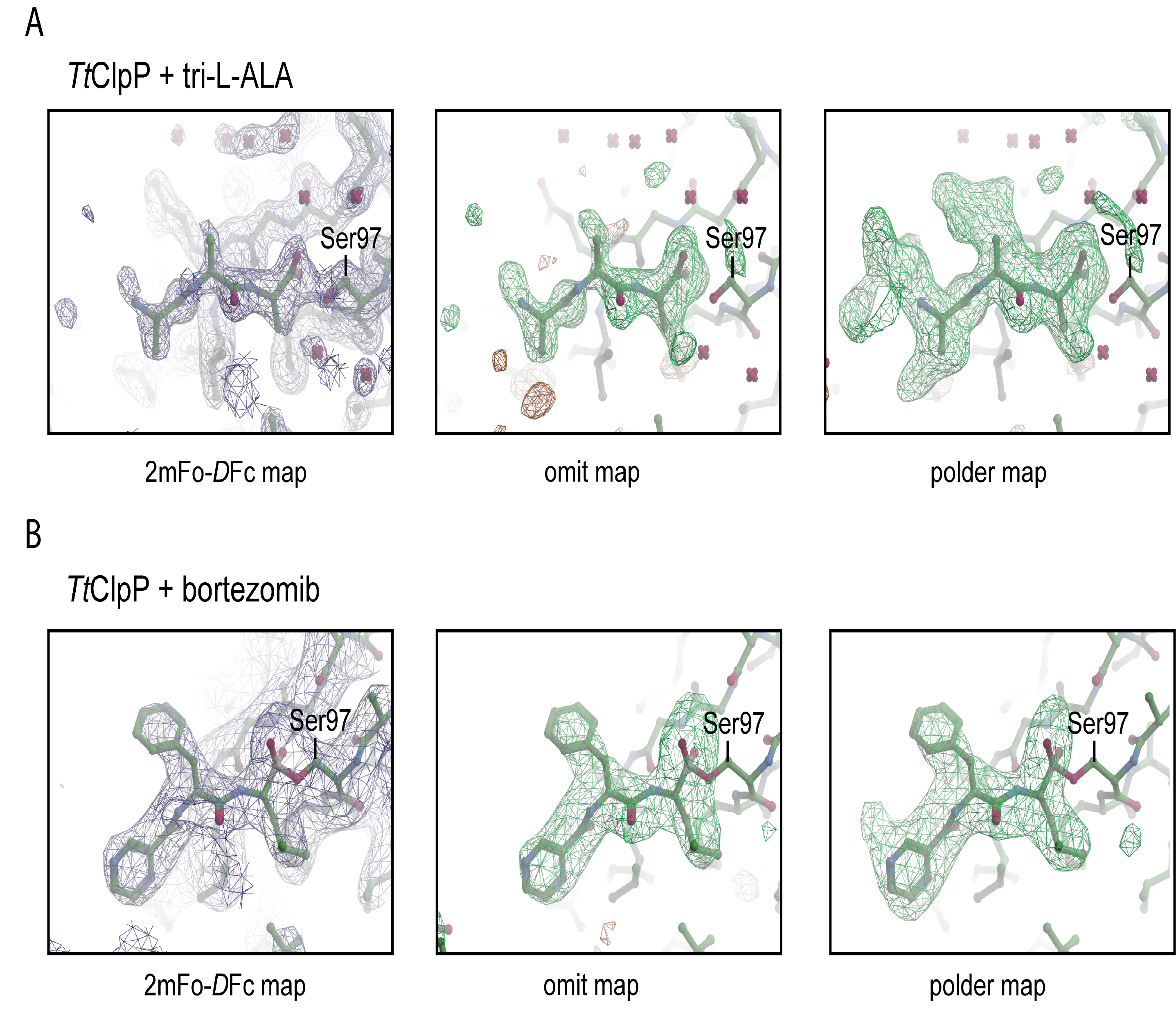
2mFo-*D*Fc, omit and polder maps of tri-L-ALA (A) and bortezomib (B) present near Ser97 in the TtClpP active site. For the omit and polder maps, mFo-*D*Fc difference density is contoured at 3σ.

**Figure Supplement 6.**
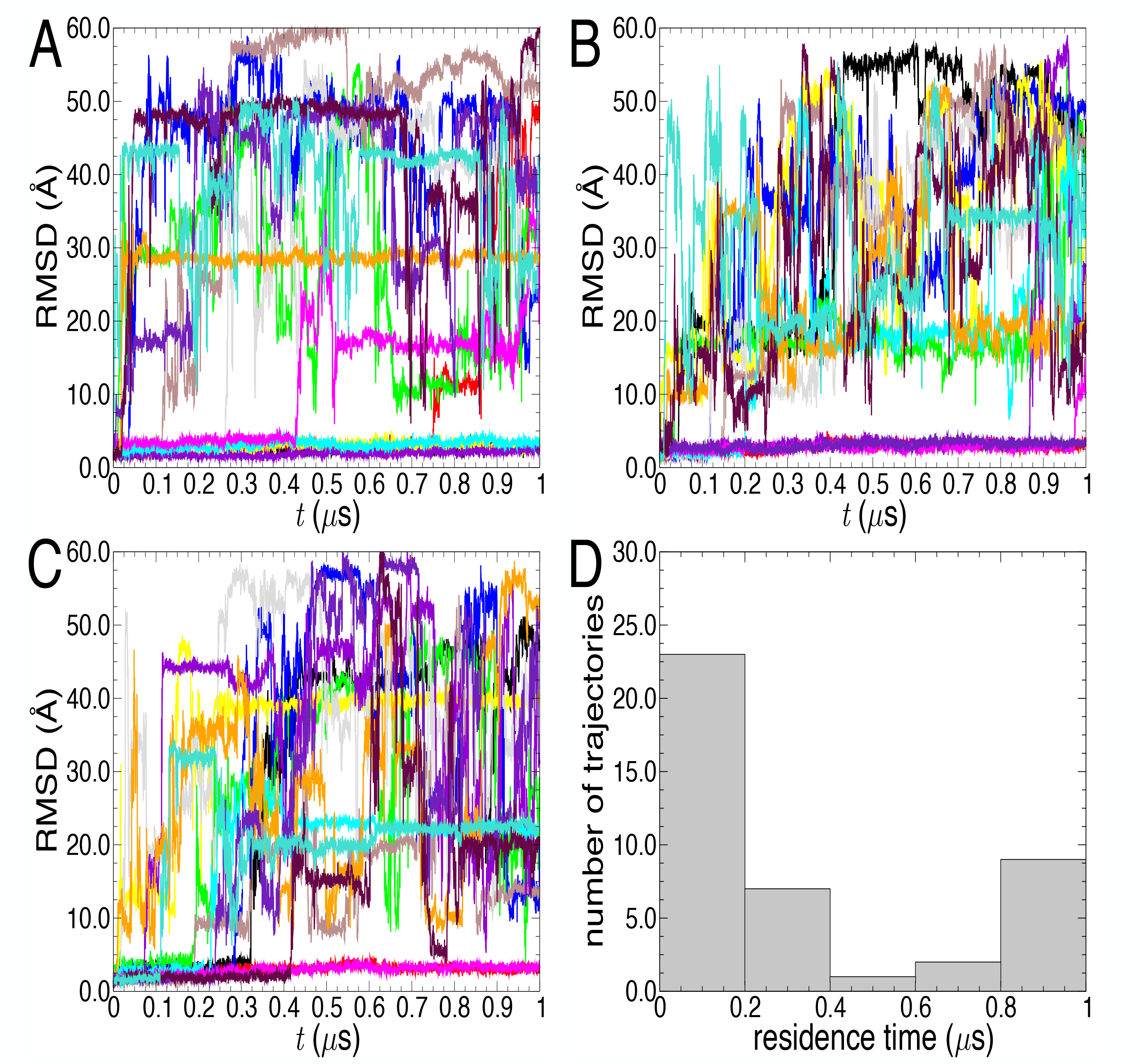
Distance root-mean-square deviation of the 14 alanine tripeptide ligands with respect to their initial bound state in ClpP, for three independent, 1-µs long molecular dynamics simulations (A-C). Profiles plateauing about 0 reflect a bound state in the initial catalytic site. Distribution of the alanine tripeptide ligands as a function of their residence time in the binding pockets (D). Out of the three simulations × 14 substrates, nine remain continuously associated to their designated catalytic site (D).

**Figure Supplement 7.**
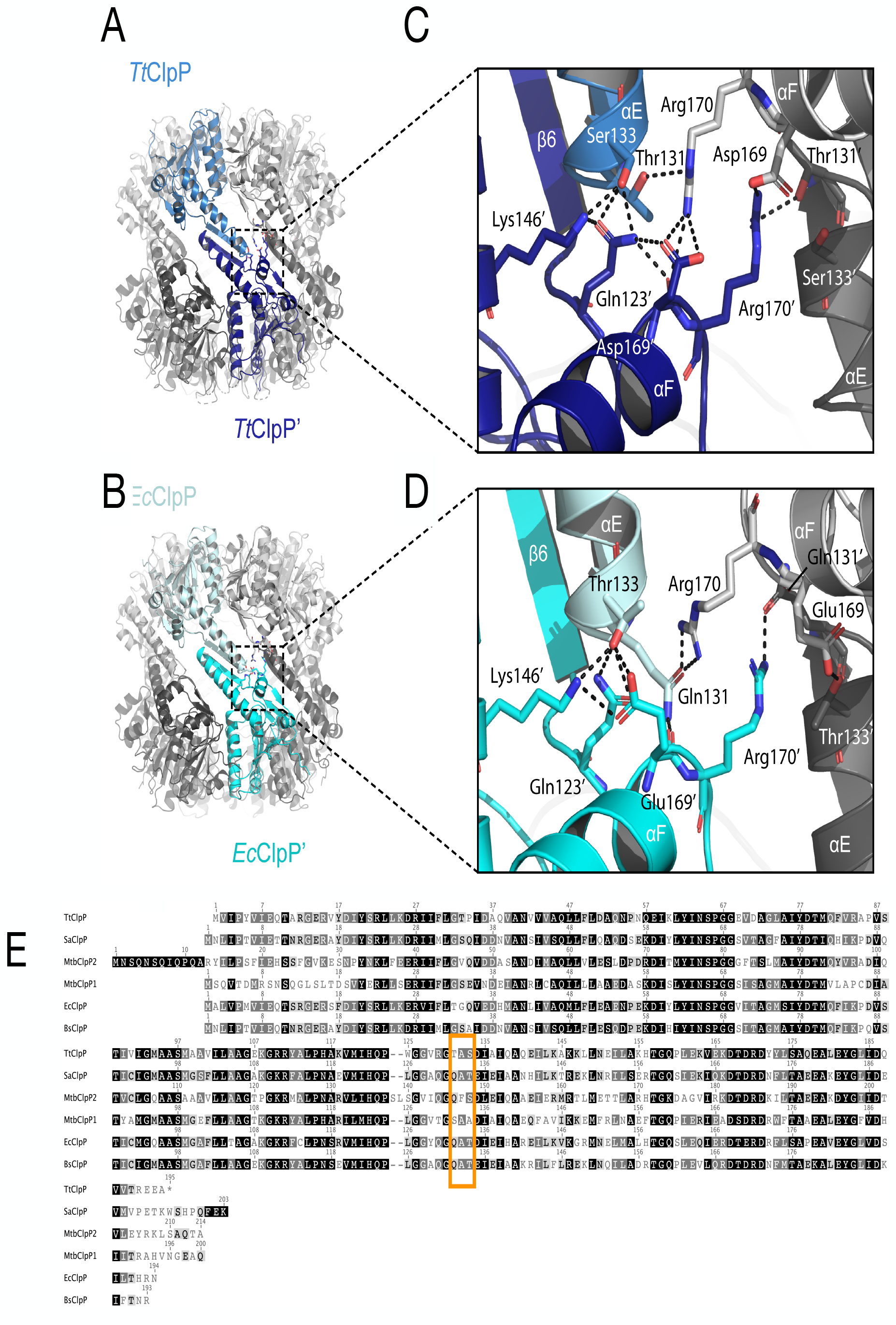
Analysis of structural differences between *Tt*ClpP and *Ec*ClpP. Although the *Tt*ClpP tetradecameric structure (A) is similar to the previously determined structures of *Ec*ClpP in the extended conformation (PDB_ID: 1 TYF) (B) differences are found at the tip of the turn that follow helix αE close to β-strand 6 in the handle domain. Residues involved in inter and/or intra-ring contacts are shown as sticks. Note the weaker hydrogen-bonding pattern formed by Thr131 in *Tt*ClpP (C) versus Gln131 in *Ec*ClpP (D). Indeed, Gln131 is conserved but replaced by Thr in *Tt*ClpP as well as *Mtb*ClpP1 (E).

**Table S1:**
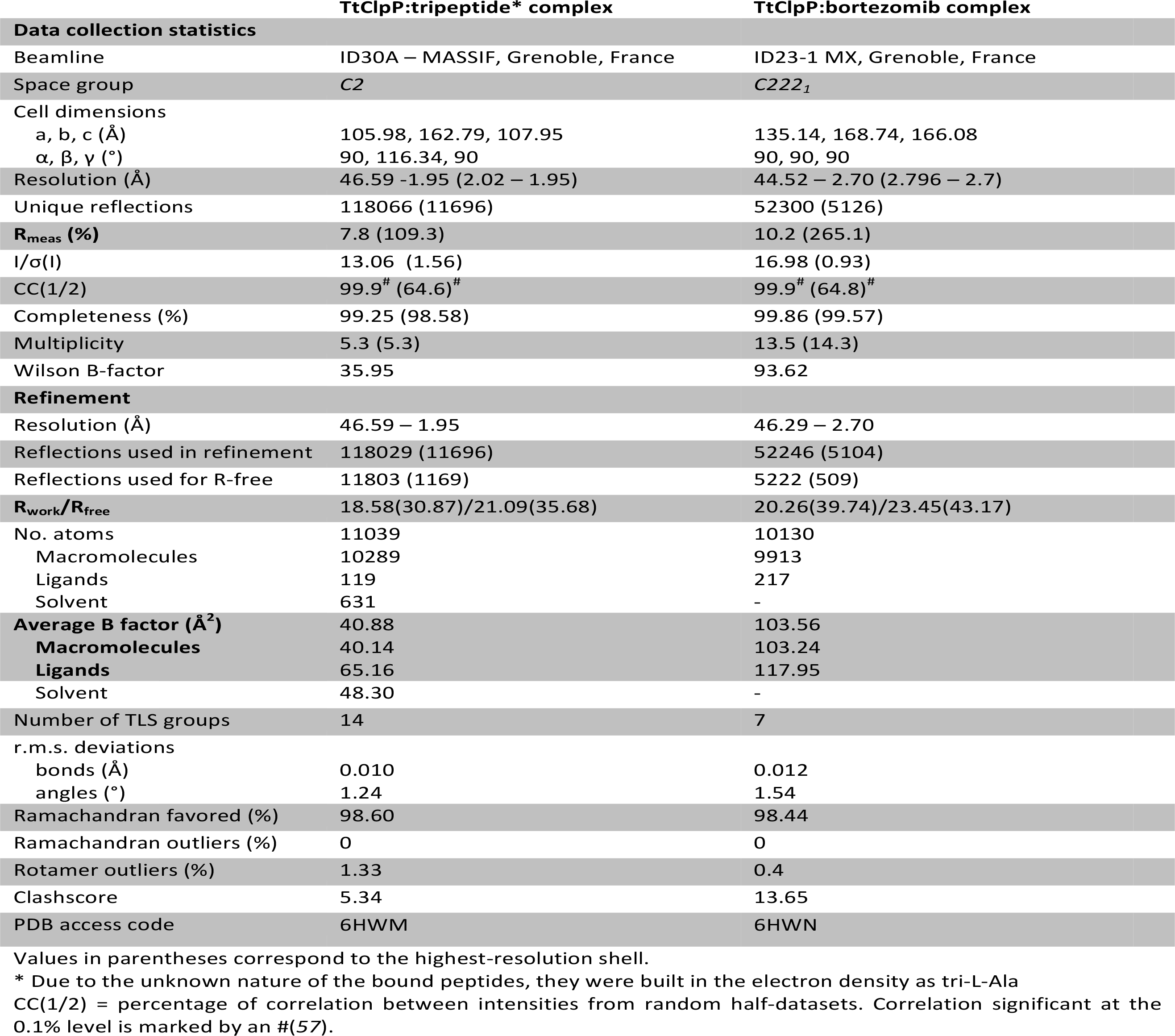
Crystallographic data collection and refinement statistics.

|  | TtClpP:tripeptide* complex | TtClpP:bortezomib complex |
| --- | --- | --- |
| **Data collection statistics** |  |  |
| Beamline | ID30A – MASSIF, Grenoble, France | ID23-1 MX, Grenoble, France |
| Space group | C2 | C222<sub>1</sub> |
| Cell dimensions |  |  |
| a, b, c (Å) | 105.98, 162.79, 107.95 | 135.14, 168.74, 166.08 |
| α, β, γ (°) | 90, 116.34, 90 | 90, 90, 90 |
| Resolution (Å) | 46.59 -1.95 (2.02 – 1.95) | 44.52 – 2.70 (2.796 – 2.7) |
| Unique reflections | 118066 (11696) | 52300 (5126) |
| R<sub>meas</sub> (%) | 7.8 (109.3) | 10.2 (265.1) |
| I/σ(I) | 13.06 (1.56) | 16.98 (0.93) |
| CC(1/2) | 99.9<sup>#</sup> (64.6)<sup>#</sup> | 99.9<sup>#</sup> (64.8)<sup>#</sup> |
| Completeness (%) | 99.25 (98.58) | 99.86 (99.57) |
| Multiplicity | 5.3 (5.3) | 13.5 (14.3) |
| Wilson B-factor | 35.95 | 93.62 |
| **Refinement** |  |  |
| Resolution (Å) | 46.59 – 1.95 | 46.29 – 2.70 |
| Reflections used in refinement | 118029 (11696) | 52246 (5104) |
| Reflections used for R-free | 11803 (1169) | 5222 (509) |
| R<sub>work</sub>/R<sub>free</sub> | 18.58(30.87)/21.09(35.68) | 20.26(39.74)/23.45(43.17) |
| No. atoms | 11039 | 10130 |
| Macromolecules | 10289 | 9913 |
| Ligands | 119 | 217 |
| Solvent | 631 | - |
| **Average B factor (Å<sup>2</sup>)** | 40.88 | 103.56 |
| **Macromolecules** | 40.14 | 103.24 |
| **Ligands** | 65.16 | 117.95 |
| Solvent | 48.30 | - |
| Number of TLS groups | 14 | 7 |
| r.m.s. deviations |  |  |
| bonds (Å) | 0.010 | 0.012 |
| angles (°) | 1.24 | 1.54 |
| Ramachandran favored (%) | 98.60 | 98.44 |
| Ramachandran outliers (%) | 0 | 0 |
| Rotamer outliers (%) | 1.33 | 0.4 |
| Clashscore | 5.34 | 13.65 |
| PDB access code | 6HWM | 6HWN |

Values in parentheses correspond to the highest-resolution shell.
\* Due to the unknown nature of the bound peptides, they were built in the electron density as tri-L-Ala
CC(1/2) = percentage of correlation between intensities from random half-datasets. Correlation significant at the 0.1% level is marked by an # (57).

